# Nuclear and cytoplasmic RNA exosomes and PELOTA1 prevent miRNA-induced secondary siRNA production in Arabidopsis

**DOI:** 10.1101/2021.05.31.446391

**Authors:** Maria Louisa Vigh, Axel Thieffry, Laura Arribas-Hernández, Peter Brodersen

## Abstract

Amplification of short interfering RNA (siRNAs) via RNA dependent RNA Polymerases (RdRPs) is of fundamental importance in RNA silencing. In plants, silencing by microRNAs (miRNAs) generally does not lead to engagement of RdRPs, in part thanks to an as yet poorly understood activity of the cytoplasmic exosome adaptor SKI2. Here, we show that mutation of the cytoplasmic exosome subunit RRP45B results in siRNA production very similar to what is observed in *ski2* mutants. Furthermore, loss of the nuclear exosome adaptor HEN2 leads to secondary siRNA production from miRNA targets largely distinct from those producing siRNAs in *ski2*. Importantly, mutation of the Release Factor paralogue PELOTA1 required for subunit dissociation of stalled ribosomes causes siRNA production from miRNA targets overlapping with, but distinct from, those affected in *ski2* and *rrp45b* mutants. We also show that miRNA-induced illicit secondary siRNA production correlates with miRNA levels rather than accumulation of stable 5’-cleavage fragments. We propose that stalled RNA-induced Silencing Complex (RISC) and ribosomes, but not stable target mRNA cleavage fragments released from RISC, trigger secondary siRNA production, and that the exosome limits siRNA amplification by reducing RISC dwell time on miRNA target mRNAs while PELOTA1 does so by reducing ribosome stalling.

## INTRODUCTION

Small RNA-guided repression of mature mRNA involves a simple, linear pathway. Dicer ribonucleases process RNA with double-stranded features into 20-24-nt duplexes with 2-nt 3’-overhangs, and one of the two strands stably associates with an Argonaute (AGO) protein to form a mature, minimal RNA-induced Silencing Complex (RISC). RISC then uses the base pairing specificity of the small RNA to find complementary targets, and it may use the endonuclease activity of AGO to cleave target RNA, the process referred to as slicing (1,2). The employment of this simple pathway gives rise to repression that can be relieved by degradation or arrest of biogenesis of the small RNA. In plants, nematodes and many fungi, an additional positive feedback module is part of small RNA-mediated gene regulation, as RNA-dependent RNA Polymerases (RdRPs) may be recruited to mRNAs targeted by RISC loaded with a primary small RNA species (3,4). In nematodes, direct RdRP products guide target RNA silencing (5), whereas in plants, RdRP activity gives rise to synthesis of target-templated double-stranded RNA that serves as substrate for DICER-LIKE proteins to generate more small interfering RNAs (siRNAs) complementary to the target (6-8). These amplified siRNAs are referred to as secondary siRNAs, and reiterative rounds of amplification may then generate tertiary siRNAs, quaternary siRNAs etc. Regardless of the exact mechanism of secondary small RNA biogenesis, the decision to employ small RNA amplification represents a major checkpoint of genetic control, because it determines the resulting type of regulation. In plants, three fundamentally different outcomes are possible dependent on whether and how small RNA amplification is employed: First, in the total absence of amplification, silencing is reversible and dependent on the continued production of the primary small RNA species. In addition, in the case of miRNAs, the regulation is typically cell-autonomous or restricted to spreading over a few cells by cell-to-cell movement (9-12), although important examples of miRNA action over long distances via phloem transport have also been reported (13,14). Second, if a single round of secondary siRNAs is produced, gene regulation remains reversible and dependent on the primary trigger (15), but the regulatory RNAs move efficiently from cell to cell (12). This phenomenon has profound biological ramifications as it may result in formation of concentration gradients and consequent action of siRNAs as morphogens during development (16,17), and in vesicle-mediated siRNA-transfer into fungal and oomycete pathogens as part of immune responses (18,19). The typical example of such small RNAs resulting from single amplification rounds is *trans-*acting siRNAs (tasiRNAs), whose accumulation pattern in phase with a miRNA-directed cleavage site is proof of single-round amplification (3,7,8). Third, if reiterative amplification is produced, silencing is effectively irreversible as it becomes independent of the primary trigger, and maintains the target silenced. Endogenous transcripts are typically subjected to the first two reversible outcomes of silencing, while the third option is used to silence foreign RNA, for example transgene, transposon or viral RNA (20-25). Thus, the employment of the RdRP amplification module represents a central decision in gene regulation and is tightly linked to the fundamental problem of distinguishing self-from non-self RNA.

Although the RdRP RDR6 was the first factor required for small RNA-guided transgene silencing to be identified in plants (20,21), only few fundamental questions regarding its engagement in production of double-stranded RNA substrates for Dicers have been answered. Careful examination of its substrate preferences *in vitro* showed that RDR6 activity is nearly completely inhibited by a 3’ stretch of more than 8 adenosines (26), thus providing a simple explanation for how functional, polyadenylated mRNAs escape use as an RDR6 substrate. On the other hand, while it is now clear that 22-nt, but not 21-nt, miRNAs tend to trigger RDR6-dependent, secondary siRNA production (27-29), there is no clear explanation for why the two different size classes of miRNAs exhibit this fundamentally different behaviour. The most fruitful way to put this question may be to ask which mechanisms allow cells to avoid secondary siRNA production in target interactions involving 21-nt miRNAs, and which properties of 22-nt miRNAs cause those protective mechanisms to be overridden.

Studies of small RNA-production in mutants with lesions in key factors of mRNA decay, such as core components or activators of decapping, and the 5’-3’ exoribonuclease XRN4 (30-35), have shown that a wide range of mRNAs become sources of RDR6-dependent siRNAs in these backgrounds. Such results are generally interpreted as support for the model that aberrant RNA may initiate RDR6-dependent siRNA production. Because the sets of mRNAs giving rise to siRNA production in these mutants are generally not significantly enriched in miRNA targets, these studies do not provide useful answers to the problem of how endogenous miRNA targets escape RDR6-dependent siRNA formation.

Our previous work demonstrated that one mechanism of avoiding illicit RDR6-dependent secondary siRNAs induced by 21-nt miRNAs in Arabidopsis involves the DExH-type RNA helicase SUPERKILLER2 (SKI2), because a limited number of mRNAs strongly enriched in known miRNA targets accumulates secondary siRNAs mapping close to miRNA binding sites in *ski2* mutants (36). SKI2 is part of the heterotetrameric, cytoplasmic SKI2-3-8 complex that acts as an adaptor for the RNA exosome (37-39), the major eukaryotic 3’-5’-exoribonuclease complex (40,41). Consistent with this biochemical function, mutation of each of *SKI2, SKI3* and *SKI8* results in accumulation of stable RISC 5’-cleavage fragments from several miRNA targets (36). Cleaved mRNAs without stop codons, resulting in stalled ribosomes with empty A-sites constitute an important class of substrates of the SKI2-SKI3-SKI8-exosome pathway (42,43). In animals, this so-called non-stop decay (NSD) pathway depends on recognition of stalled ribosomes at or close to mRNA 3’-ends by the Release Factor (RF)-like proteins Pelota (in yeast referred to as Dom34) and Hbs1 (44,45) that cause ribosome subunit dissociation and release of intact peptidyl-tRNA to achieve ribosome recycling (46). 5’-cleavage fragments generated through miRNA-guided cleavage are generally NSD substrates, because most miRNA binding sites occur in open reading frames in plants (47-49). Indeed, studies of the *Arabidopsis* orthologues of Pelota and Hbs1 showed that inactivation of these NSD factors also results in accumulation of stable RISC 5’-cleavage fragments, similar to mutation of *SKI2*, with plant PELOTA being indispensable and HBS1 playing an accessory role (50). The relative importance of PELOTA and HBS1 is similar to what was previously observed for ribosome subunit dissociation from stalled elongation complexes in other eukaryotic systems (51). The *Arabidopsis* studies also revealed that plants encode two PELOTA homologues, PEL1 and PEL2, and that PEL1 is the functional homologue of metazoan Pelota, while PEL2 appears to act as a dominant negative NSD factor, perhaps through its intact HBS1-binding activity (52).

Given the lack of a 3’ poly(A)-tail of 5’-cleavage fragments and the inhibitory activity of oligo(A) tracts on RDR6 activity, combined with genetic evidence for the implication of aberrant RNA in siRNA production, it is tempting to speculate that mutation of *SKI2* indirectly allows secondary siRNA formation via action of RDR6 on stable 5’-cleavage fragments. This seemingly straight-forward interpretation is in conflict with several observations, however. First, for some miRNA targets, stable 5’-cleavage fragments, but no secondary siRNAs, can be detected in *ski2* mutants (36). Second, for other miRNA targets, secondary siRNAs accumulate exclusively 3’ to the cleavage site despite clear accumulation of 5’-cleavage fragments, precluding action of RDR6 on stable 5’-cleavage fragments as the general mechanism of illicit secondary siRNA formation in *ski2* mutants (36). Thus, although SKI2 is part of one of the molecular mechanisms that guards against siRNA amplification induced by 21-nt miRNAs, it is unclear how it does so. A key prerequisite for a better understanding of this important problem is to clarify whether the two effects of SKI2 rely on the same biochemical activity – acceleration of exosome-mediated degradation of 5’-cleavage fragments – or whether SKI2 uses a separate activity to ensure avoidance of secondary siRNA production. We reasoned that if the RNA decay activity of the SKI2/exosome pathway is implicated in avoiding miRNA-triggered secondary siRNAs, mutants in all components of this pathway should show defects in limiting secondary siRNA production similar to *ski2* mutants. On the other hand, this defect should be specific to *ski2* mutants if a function of the protein distinct from stimulation of exosome-coupled RNA decay limits secondary siRNA production.

We show here that mutants in *SKI3* and in the cytoplasmic exosome subunit *RRP45B* have defects in limitation of secondary siRNA production very similar to *ski2* mutants, and that a hypomorphic mutant in the gene encoding the core exosome subunit RRP4 has more widespread defects in limiting production of illicit siRNAs. Mutants in *RRP45B* also have defects in degradation of RISC 5’-cleavage fragments, thus indicating that SKI2-mediated exosomal decay of 5’-cleavage fragments is necessary for avoidance of miRNA-triggered secondary siRNAs. Furthermore, inactivation of the nuclear exosome cofactor *HEN2* leads to secondary siRNA generation from a subset of miRNA targets largely distinct from those affected by SKI2, SKI3, and RRP45B, perhaps suggesting that some miRNA targets may undergo RISC-mediated cleavage in the nucleus. We also show that the expression level of 21-nt miRNAs, not the accumulation of their RISC-mediated cleavage fragments, correlates with their ability to trigger secondary siRNAs in *rrp4* mutants. We discuss these results in the light of our observation in the present work that mutation of *PEL1* leads to miRNA-induced siRNA amplification similar, but not identical, to that observed in *ski2, ski3* and *rrp45b*, and of recent evidence for the importance of ribosome stalling for siRNA production (53).

## MATERIAL AND METHODS

### Plant material and growth conditions

All mutants used in this study are of the ecotype Columbia (Col-0). T-DNA insertion mutants in *SKI2* (AT3G49690: *ski2-5* (SALK_118529)), *SKI3* (AT1G76630: *ski3-5* (GK_140B07)), *SKI8* (AT4G29830: *ski8-1* (SALK_083364)), *HEN2* (AT2G06990: *hen2-4* (SALK_091606), *hen2-5* (GK_313G10_015818)), *PEL1* (AT4G27650: *pel1-1* (SAIL_881_B10)), RRP45A (AT3G12990: *rrp45a* (GK_655_D02)), *RRP45B* (AT3G60500: *cer7-3* (SAIL_747_B08) and *cer7-3*/*rdr6-12*)) and *SOP1* (AT1G21580: *sop1-*5 (SALK_019457) have been described previously (36,50,54-58), as has the *rrp4-2/sop2* point mutant allele (58). *pel1-2* (GK_537F08) was identified from the GABI-KAT collection of T-DNA insertion mutants (59). All T-DNA mutants except *cer7-3, cer7-3*/*rdr6-12, hen2-4, hen2-5* and *sop1-5* were obtained from the Nottingham Arabidopsis Stock Centre. *cer7-3* and *cer7-3*/*rdr6-12* mutant seeds were a gift from Ljerka Kunst, *hen2-4* mutant seeds were a gift from Dominique Gagliardi, while *hen2-5, sop1-5* and *rrp4-2* mutant seeds were a gift from Kian Hématy. All experiments were carried out with inflorescence tissue. To obtain this tissue, seedlings were germinated on plates containing 1xMurashige & Skoog medium, supplemented with 1% /w/v) sucrose and 0.8% (w/v) agar, transferred to soil ((Plugg/Såjord [seedcompost];; SW Horto)) 10 days after germination (DAG) and grown in a Percival growth chamber for an additional four weeks under the following conditions: 16 h light (Master TL-D 36W/840 and 18W/840 bulbs (Philips);; 130 mmol m^-2^ s^-1^, 21 °C, 60% relative humidity)/ 8 h darkness (16°C, 60% relative humidity). Inflorescences were collected and snap-frozen in liquid nitrogen.

### Mutant genotyping

Each T-DNA insertion mutant was genotyped with two PCR reactions: a PCR reaction with a set of primers flanking the insertion site for wild type allele detection and a PCR reaction with an outward left border primer and one of the two flanking primers for T-DNA allele detection. Genotyping of *rrp4-2* was carried out with a single PCR reaction with primers spanning the point mutation followed by restriction analysis (Eco47I). Genotyping of *rdr6-12* was carried out as described in (60). For construction of the *hen2-4/ski2-5* double mutant, F2 populations generated by self-pollination of F1 plants obtained from *hen2-4* crossed to *ski2-5* were genotyped as described above to find double homozygous mutants. All primer sequences used in this study are listed in **Supplementary Table S1**.

### RNA extraction

Total RNA from inflorescences was extracted with TRI Reagent (Invitrogen) according to the manufacturer’s instructions. The RNA was dissolved in either 50% formamide for northern blots or in sterilized water for small RNA libraries.

### Northern blotting of mRNA cleavage fragments

20 μg of purified total RNA was loaded onto a 1% denaturing agarose gel and separated for 3 h at 120V. The RNA was blotted to an Amersham Hybond-NX membrane (GE Healthcare Life Sciences) followed by UV-crosslinking (254 nm). Two gels with the same batch of extracted RNA were prepared in parallel. After pre-hybridization in PerfectHyb^™^ Plus Hybridization buffer (Sigma) for 1 h at 65 °C, the two blots were hybridized O/N at 65 °C to each their radioactively labeled 3’ CF probe (AGO1 3’ CF and CSD2 3’ CF probes labelled with Prime-a-gene labeling kit from Promega). The membranes were washed 3 times in 2xSSC (0.3 M NaCl, 30 mM sodium citrate), 0.1% SDS at 65°C and developed by phosphorimaging. The blots were subsequently stripped with boiling 0.1% SDS and rehybridized with each their radioactively labelled 5’ CF probes. Sequences for primers used for amplification of probe templates of AGO1 and CSD2 cleavage fragments are listed in **Supplementary Table S1**.

### Organisation of small RNA sequencing experiments

This study includes analysis of raw small RNA sequencing data from three independent sets of experiments, referred to as experiments A, B and C. sRNA sequencing data from experiment A (Col-0 WT, *ski2-5, ski3-5, rrp45b*) is used in **Figure 2**. sRNA sequencing data from experiment B (Col-0 WT, *ski2-5, pel1-1, pel1-2, hen2-5, rrp4, rrp45a*) is used in **Figure 3-7**. sRNA sequencing data from experiment C (Col-0 WT, *ski2-5, hen2-5, sop1-5*) is used in **Figure S4**. Experiment A includes 2 biological replicates of each genotype, while experiment B and C were performed using three biological replicates for each genotype, except for the genotype *pel1-1*, which is represented in experiment B with one biological sample due to loss of tissue during sampling. Each biological replicate represents inflorescences collected from four individual plants of the same genotype grown in the same pot.

### Construction of libraries for small RNA sequencing

1 μg of total inflorescence RNA was used as input in each library. Libraries were generated using the NEBNext Small RNA Library Prep Sets (New England Biolabs) according to the manufacturer’s instructions. The indexed cDNA libraries were size selected on a 6% polyacrylamide gel as described in the NEBNext protocol. All size-selected libraries were analyzed using an Agilent Bioanalyzer, quantified with Qubit measurements and single-end sequenced on an Illumina Nextseq 500 with SE75_HI chemistry. 1% of spike-in PhiX was also loaded on the flow cells in each of the three runs.

### Analysis of small RNA reads

#### Mapping

The adaptor sequence, AGATCGGAAGAGCACACGTCTGAACTCCAGTCAC, was trimmed from raw reads with cutadapt v.2.4 (61). Reads were mapped with STAR_2.6.0a (62) to TAIR10 with the following parameters: *outFilterMultimapNmax 20, alignIntronMax 1, outFilterMismatchNmax 1, outFilterMismatchNoverLmax 0, outFilterScoreMinOverLread 0, outFilterMatchNminOverLread 0, outFilterMatchNmin 15, alignSJDBoverhangMin 100*. No rRNA or tRNA reads were removed prior to mapping. Mapped reads were quantified with featureCounts from subread-1.6.3 (63) with an Araport11 transcriptome reference file custom modified to also include intergenic regions.

#### Initial quality controls

Reads were not quality controlled before mapping as FastQC on small reads is not recommended. On the other hand, the samples were validated based on principal component analyses and distance plot matrices prior to the differential expression analysis. Both PCA and distance plots were based on vst-transformed normalized reads (RPM) (64). One of the *ski2-5* replicates in experiment A was removed from further downstream analysis as it was a clear outlier in the PCA and distance plots (**Supplemental Figure 1A,B**). Using same argument, the sole biological replicate of *pel1-1* in experiment B was included for further downstream differential expression analysis as it clusters close to all three replicates of *pel1-2* (**Supplemental Figure 1C,D**). The data resulting from small RNA sequencing of libraries from experiment C was validated prior to differential expression analysis in the same manner, but no sample outliers were found (**Supplemental Figure 1E,F**).

#### Differential expression analysis

The package DESeq2 (64) was used for the differential expression analysis. Normalized reads were estimated for size factors (65) and a generalized linear model was used. The DESeq2 results were extracted with manual contrasting of mutant vs. WT. A gene was considered to have differential expressed small RNA levels compared to WT if the padj. < 0.05. All differentially expressed genes from the three DESeq2 analyses are listed in the supplemental tables (**Supplemental Table S2-S4**). Genes were categorized as known miRNA targets based on a list of experimentally validated targets (**Supplemental Table S5**).

#### Calculating absolute distances of small RNA reads to the cleavage sites

Genomic ranges of known miRNA targets were extracted from BigWig files and bw counts on each genomic position were normalized to the total amount of mapped reads and a mean was calculated by accounting for sample replicates. The genomic positions of a miRNA target and the 1 bp genomic position of its corresponding cleavage site were converted to transcriptomic positions using the ensembldb package (66).

#### Analysis of phasing

The first nucleotide adjacent to a known cleavage site (CS) of a miRNA target was designated as nucleotide (nt) 1 on both 5’ and 3’ CFs. Afterwards, the CFs were split into 21-nt intervals (**Figure 6A**). An siRNA was counted to belong to phase 1 if it mapped to the target transcript in any 21-nt interval arising from the CF being split from nt position 1 (e.g. 1-21, 22-42, 43-36 etc.). Only perfectly matching 21-nt long reads (cigar string = 21M) were included in the phasing analysis. The siRNAs were considered to be phased if the majority of reads fall into phase 1. The 2 nt displacement of the minus strand was not taken into consideration in the code and therefore a clear division of reads mapping to either the minus or plus strand is visible in phase 1 and 2 for the Tas1C example (**Figure 6B**).

#### Analysis of trimming and tailing

The trimming and tailing analysis of sRNA reads mapping to the extremity of the 5’ CF in *PHB* and *TCP2* was performed by filtering the BAM files on an approximately 10 nt match in sequence – 10 nt from the cleavage site (**Figure 6C**). Whether a read was trimmed or tailed and how many nucleotides were missing from or added to the end of the original CF end was computed on the basis of the sequence and cigar string.

All data analysis of the mapped and quantified reads were performed in R https://www.r-project.org/ and plots were generated with ggplot2 https://ggplot2.tidyverse.org/. All R code is available at the following Github repository: https://github.com/MariaLouisaVigh/ExosomePelota.

## RESULTS

### Inactivation of the cytoplasmic exosome results in accumulation of RISC 5’-cleavage fragments

To answer the question of whether the RNA exosome mediates the degradation of RISC 5’-cleavage fragments that we previously showed to depend on the components of the SKI complex, SKI2, SKI3, SKI8, we performed northern blot analysis of RNA prepared from *rrp45b* mutants (also known as *cer7-3*). *RRP45B* is one of two genes encoding the core exosome subunit RRP45 in Arabidopsis, and in wild type plants, RRP45B is part of cytoplasmic exosomes while the isoform, RRP45A, is part of nuclear exosomes (55,58). These analyses showed that the *rrp45b* mutant exhibits stabilization of 5’-cleavage fragments of *AGO1*, targeted by miR168, and of *CSD2*, targeted by miR398 (**Figure 1A,B**), similar to what is observed in *ski2, ski3* and *ski8* mutants. In addition, as seen previously for *ski2-4* mutants (36), mutation of *RDR6* did not affect the level of 5’-cleavage fragments detected in *rrp45b* mutants (**Figure 1A,B**). We conclude from these observations that the degradation of those RISC 5’-cleavage fragments proceeds via a canonical SKI2-3-8-exosome dependent pathway.

**Figure 1.**
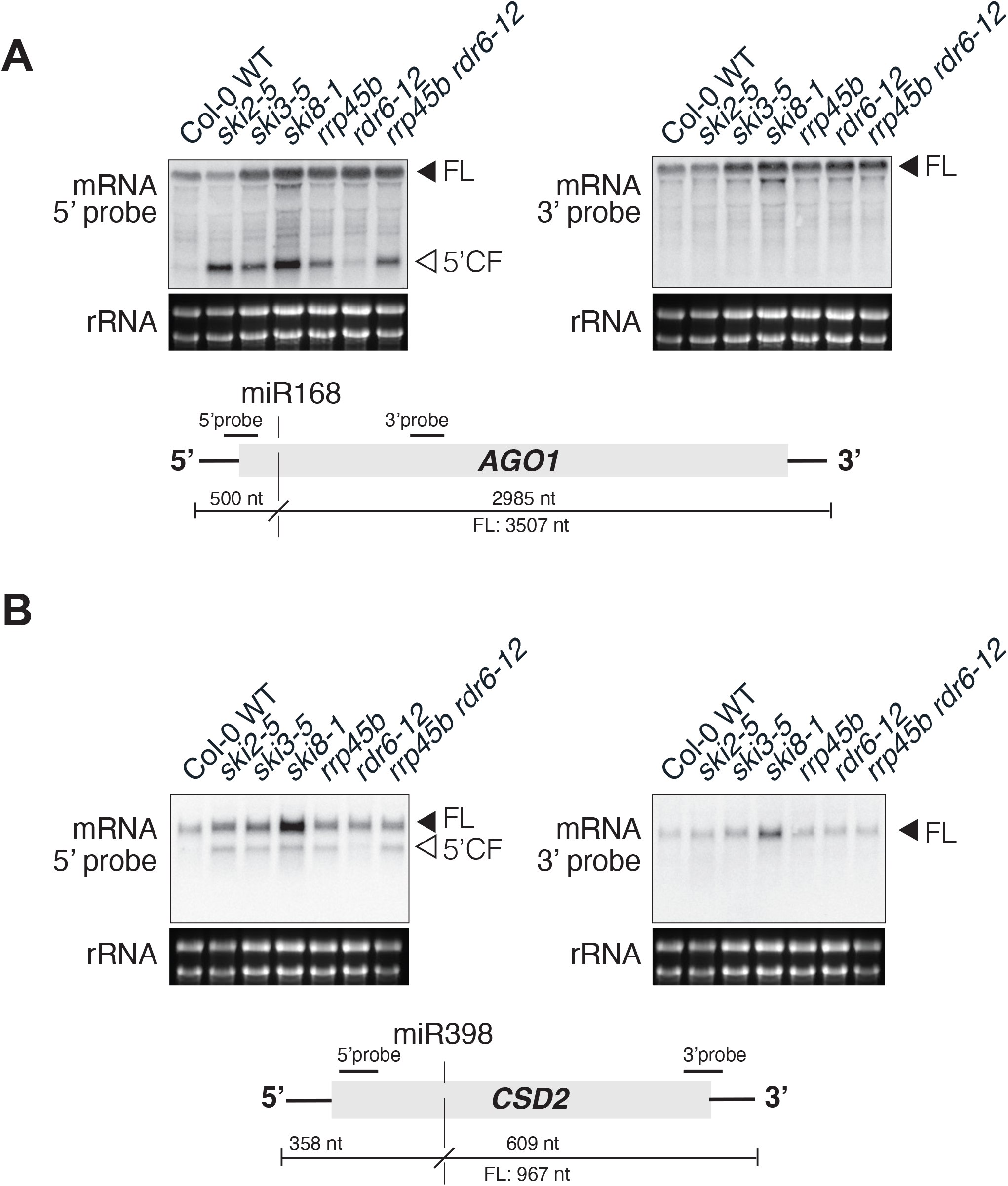
Accumulation of 5’ cleavage fragments in *rrp45b*. (**A**) Northern blot analyses of 20 µg total RNA from inflorescences. Radiolabeled probes specific to either the 5’ or the 3’ cleavage fragment of AGO1 were hybridized consecutively to the same membrane. Arrows indicate the full length (FL) and the 5’ cleavage fragment (CF). Localization of the probes relative to the miR168 cleavage site (vertical dashed line) and expected cleavage fragment sizes are illustrated below the blots. (**B**) Same analysis as in (**A**) carried out with the miR398 target CSD2.

### *rrp45b* and *ski3* mutants exhibit defective limitation of secondary siRNAs similar to ski2

We next tested whether RRP45B and SKI3 are also necessary for avoidance of miRNA-induced secondary siRNA production, similar to SKI2. Small RNA-seq with RNA extracted from *ski2, ski3* and *rrp45b* mutants showed that miRNA targets were enriched among mRNAs producing ectopic siRNAs, and that very similar sets of miRNA targets produced secondary siRNAs in the three mutants (**Figure 2A,B**). Moreover, ectopic siRNAs mapped close to miRNA-guided cleavage sites (**Figure 2C**), as described previously for *ski2-4* mutants (36). The abundance of the trigger miRNAs themselves was unchanged compared to wild type in all of the mutants (**Supplemental Figure 2A**), excluding increased trigger miRNA levels as a possible explanation for the increased siRNA production. We also verified that for all three mutants, no secondary siRNAs mapping to CSD2 were detectable (**Figure 2C**), despite the clear stabilization of CSD2 5’-cleavage fragments (**Figure 1B**). Since knockout mutants in genes encoding two distinct components of the SKI2-3-8 complex, an ATP-binding site mutant of *SKI2* (*ski2-4*) (36), and a mutant in *RRP45B* give highly similar profiles of illicit secondary siRNAs on miRNA targets, and lead to similar stabilization of 5’-cleavage fragments, we conclude that SKI2-mediated exosome function, presumably 3’-5’ exonucleolysis of 5’-cleavage fragments, underlies its role in limitation of secondary siRNA production. We stress that this is a not a trivial result: the free, stable 5’-cleavage fragment cannot be the direct template for RDR6 in those cases in which secondary siRNAs map to the 3’-cleavage fragment, and is generally unlikely to serve as an RDR6 template because of the existence of examples such as CSD2 for which stable 5’-cleavage fragments do not give rise to siRNA populations. Thus, exosomal degradation of 5’-cleavage fragments is required to avoid siRNA production and spreading in both 5’- and 3’-directions relative to the miRNA-guided cleavage site, a point that will be treated in depth in the discussion. Consistent with our results, miRNA targets have previously been observed to be represented among siRNA-generating transcripts in mutants of the exosome or of the RST1/RIPR complex implicated in connecting cytoplasmic exosome and SKI2-3-8 complexes (33,35).

**Figure 2.**
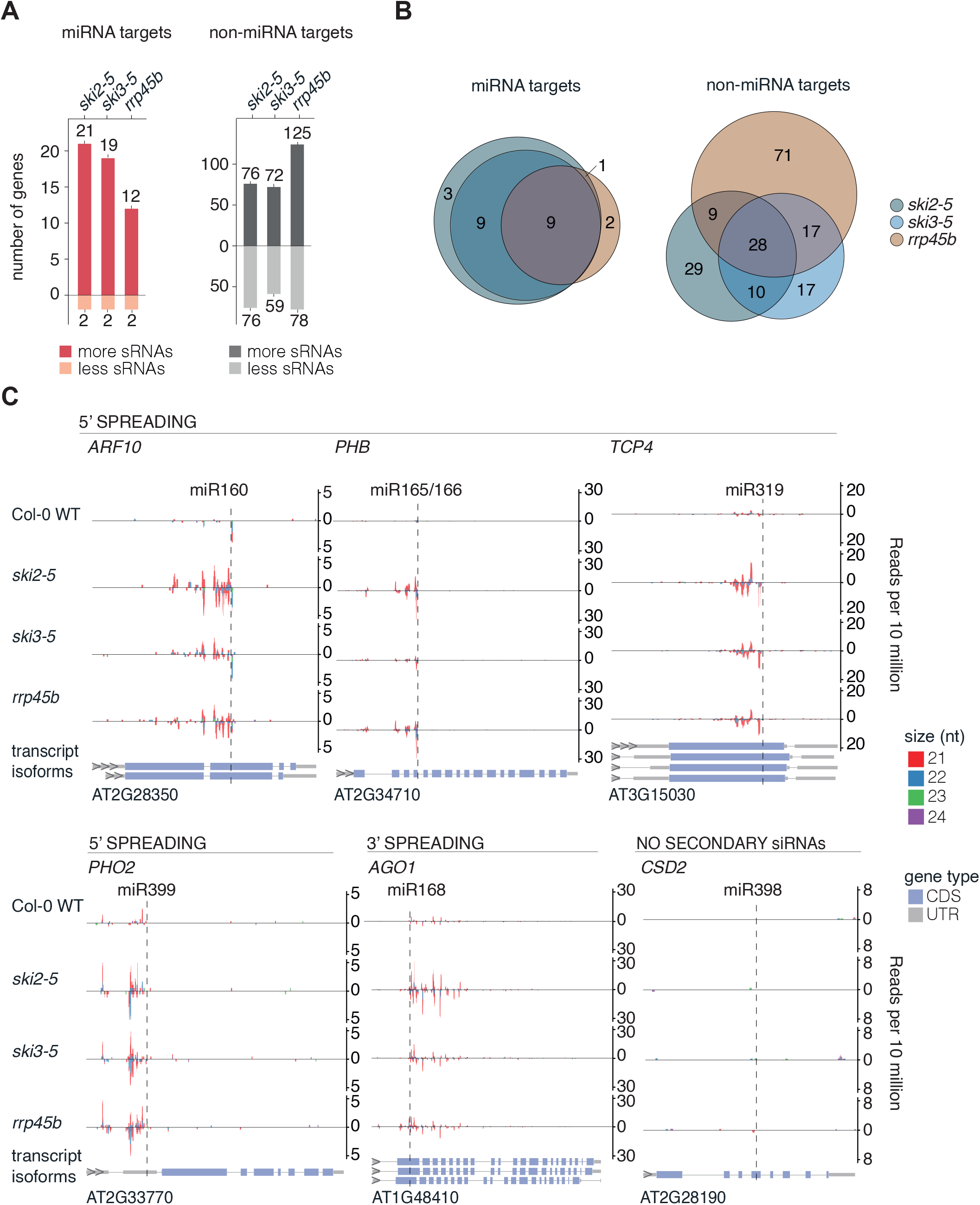
SKI3 and RRP45B limit secondary siRNA production on miRNA targets. (**A**) Bar plots depicting number of known miRNA targets (red bars) and number of mRNAs not known to be miRNA targets (non-miRNA targets, grey bars) which produce either more or less secondary siRNAs in *ski2-5, ski3-5* or *rrp45b* compared to Col-0 WT (Wald test, P < 0.05). The enrichment of miRNA targets in genes producing more siRNAs are highly significant in *ski2-5, ski3-5* and *rrp45b* (Fisher-test for *ski2-5* and *ski3-5*: P < 2.2 × 10^−16^, Fisher-test for *rrp45b*: P = 1.39 × 10^−15^). (**B**) Euler diagram of the overlap in known miRNA targets (top) and the overlap in non-miRNA targets (bottom) with ectopic siRNA accumulation in *ski2-5, ski3-5* and *rrp45b*. (**C**) Examples of siRNA accumulation on endogenous miRNA targets. Four examples with siRNAs mapping to the 5’ cleavage fragment (*ARF10, PHB, TCP4* and *PHO2*), one example of siRNAs mapping to the 3’ cleavage fragment (*AGO1*), and one example of no siRNA accumulation despite detection of a stable 5’ cleavage fragment (*CSD2*) are shown. The y-axis shows the number of small RNA reads per 10 million. Positive read counts indicate sense orientation, negative read counts indicate antisense orientation. The x-axis is the TAIR10 genomic coordinates of the target. sRNAs of sizes 21-24-nt are plotted and the sizes are distinguished with colors as indicated. Vertical dashed lines indicate miRNA-guided cleavage sites. The different transcript isoforms are illustrated for each gene with 5’- and 3’-UTRs in grey, exons in blue and introns as connecting lines. All miRNA targets are plotted in the 5’ to 3’ reading direction and their gene identifiers are noted below the transcript isoforms. Small RNA libraries prepared from two biological replicates were used for all analyses except for *ski2-*5 for which a single replicate had to be used, see Materials and Methods and **Supplemental Figure 1A,B**.

### miRNA-induced secondary siRNA production upon inactivation of *PEL1*

We next analyzed the effect of inactivation of the gene encoding PEL1. Metazoan Pelota is necessary for subunit dissociation of stalled ribosomes with an empty A-site, for example in the NSD pathway that eliminates faulty mRNAs without a stop codon (42). Consistent with this biochemical role, *pel1* mutants accumulate 5’-cleavage fragments generated by some miRNA-guided RISCs with targets in open reading frames, but not in 3’-UTRs (50). We found that *pel1* mutants (*pel1-1, pel1-2*) produced siRNAs from a limited set of mRNAs significantly enriched in known miRNA targets (**Figure 3A**), similar to *ski2-5*. The amplified siRNAs mapped adjacent to miRNA target sites in individual target mRNAs (**Figure 3B;; Supplemental Figure 3A**), consistent with a miRNA-RISC-triggered process. Importantly, the abundances of the trigger miRNAs themselves were unchanged compared to wild type in all of the mutants (**Supplemental Figure 2B**), arguing against a trivial explanation of increased RISC activity as the cause of secondary siRNA production. Furthermore, the overlap in siRNA-producing miRNA-targets between *pel1* and *ski2* mutants was significant, in contrast to the overlap in non-miRNA targets (**Figure 3C**). Nonetheless, around half of the miRNA-targets found to produce increased siRNA levels in *pel1* mutants did not do so in *ski2-5*. The opposite statement was also true, as roughly one third of the siRNA-overproducing miRNA targets found in *ski2-5* did not produce increased levels of siRNAs in either *pel1* mutant (**Figure 3C**). More generally, increased siRNA levels from miRNA targets in *ski2-5* were not correlated with those in *pel1* mutants (**Figure 3D;; Supplemental Figure 3B**), an important point that is well illustrated by inspection of individual examples. *PHO2/UBC24* targeted by miR399 is an expected example of increased secondary siRNA production specifically in *ski2-5*, but not in *pel1* mutants (**Figure 3B,D**), because the miR399 target sites are located in the 5’-UTR (**Figure 3B**), where 80S ribosomes required for recognition by PEL1 are not assembled. However, even for targets with miRNA sites in open reading frames, siRNA overproduction specific to *ski2-5* could be observed, as illustrated clearly by the examples miR168-*AGO1* mRNA and also to some extent miR160-*ARF10* (**Figure 3B,D**). *AGO1* mRNA consistently gives rise to much higher siRNA levels mapping to the 3’-cleavage fragment in mutants of the exosome and SKI complexes (**Figure 2C**), but not in any of the two independent *pel1* mutants (**Figure 3B**). Similarly, the increase in siRNAs for ARF10 in *pel1* mutants was much smaller than that observed in *ski2* mutants (**Figure 3B,D**). Conversely, miR157-*SPL2* gives rise to secondary siRNAs mapping to the 5’-cleavage fragment of SPL2 mRNA only in *pel1* mutants, and miR472 triggers more robust increases in siRNAs mapping to the 3’-cleavage fragment of *RSG2* mRNA in *pel1* than in *ski2-5* mutants (**Figure 3B, D**). These observations show that loss of PEL1 function leads to illicit secondary siRNA production from miRNA targets, similar, but not identical, to the consequence of inactivation of SKI2-3-8 and exosome complexes.

**Figure 3.**
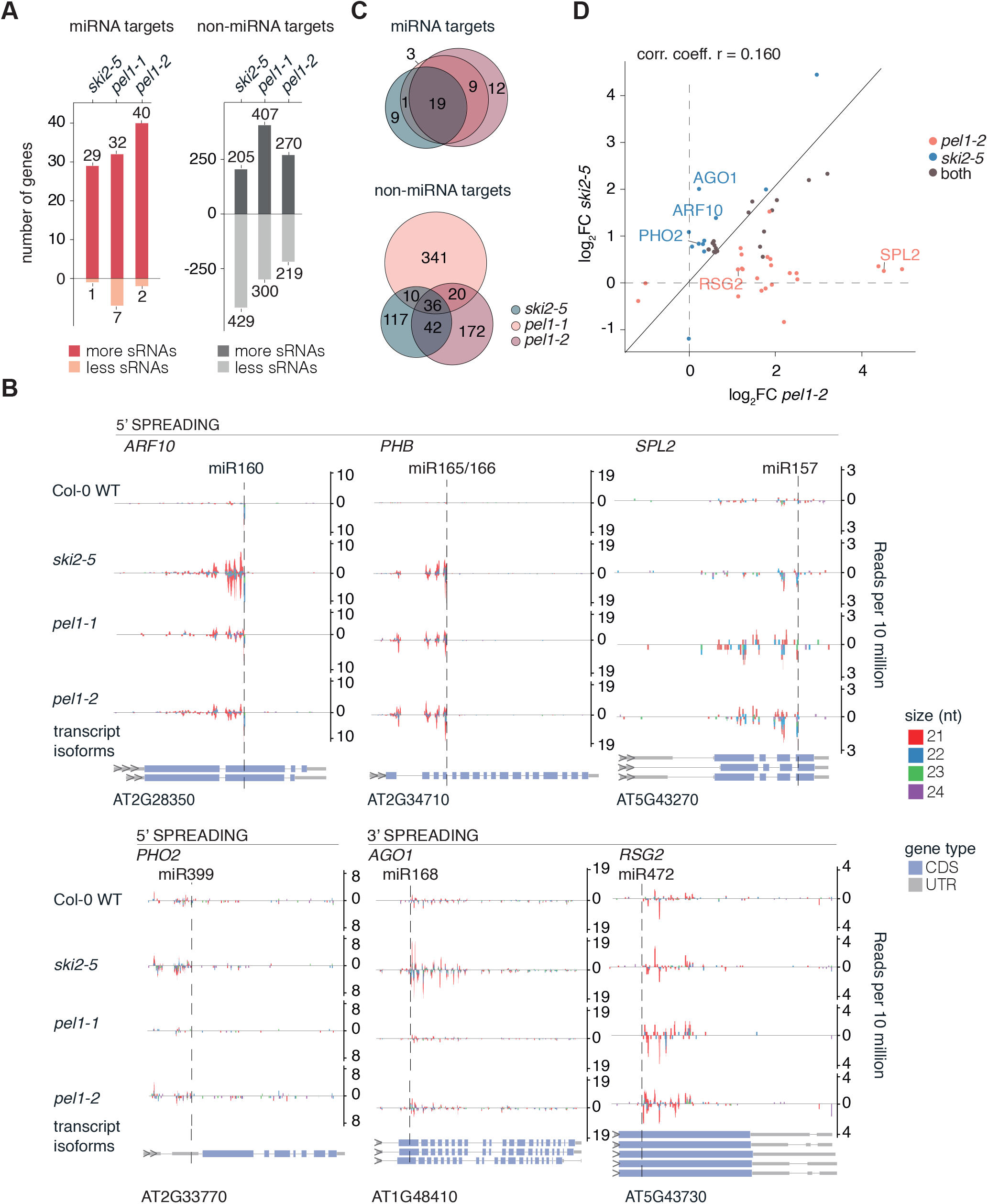
PEL1 limits secondary siRNA amplification from miRNA targets. (**A**) Bar plots depicting number of known miRNA targets (red bars) and number of mRNAs not known to be miRNA targets (non-miRNA targets, grey bars) which produce either more or less secondary siRNAs in *ski2-5, pel1-1* or *pel1-2* compared to Col-0 WT (Wald test, P < 0.05). The enrichment of miRNA targets in genes producing more siRNAs is highly significant in *ski2-5, pel1-1* and *pel1-2*, (P < 2.2 × 10^−16^ in all cases, Fisher-test). (**B**) Examples of siRNA accumulation on endogenous miRNA targets, organized as in Figure 2C. Four examples with siRNA accumulation on the 5’ CF (*ARF10, PHB, PHO2*, and *SPL2*), and two examples of accumulation on the 3’ CF (*AGO1* and *RSG2*) are shown. (**C**) Euler diagrams showing the overlaps in known miRNA targets (top) and in non-miRNA targets (bottom) with ectopic siRNA accumulation in *ski2-5, pel1-1* and *pel1-2*. The number of miRNA targets in the intersection between *ski2-5, pel1-1* and *pel1-2* is highly significant (Fisher-test: P = 1.16 × 10^−8^). (**D**) Scatter plot of log2 of the fold change of read densities (RPM(mutant)/RPM(WT)) of siRNAs (log2 FC) mapped to miRNA targets in *pel1-2* (x-axis) and *ski2-5* (y-axis). Only miRNA targets with significantly higher siRNA read counts in either mutant compared to Col-0 are included. miRNA targets shown in Figure 3B are indicated. Small RNA libraries prepared from biological triplicates were used for all analyses presented in this figure (except for *pel1-1*, which is a sample without a replicate (Supplementary Figure 1C,D). Note that the experiment reported here is fully independent from the one reported in Figure 2. Together with the fact this experiment includes more replicates and thus has more power to identify differential siRNA accumulation, this explains the different number of genes producing siRNAs detected in *ski2-5* in the experiment reported in Figure 2 and in this figure.

### The nuclear exosome broadly suppresses siRNA production

Given the importance of cytoplasmic exosome-dependent RNA decay systems for the avoidance of miRNA-triggered siRNA production, we next asked whether nuclear exosome-dependent RNA decay may also play a role. We initially analyzed mutants in two components: Knockout alleles of *HEN2* which encodes a nucleoplasmic relative of SKI2 (54), and a hypomorphic allele of *RRP4* encoding a core exosome component *(56)*, expected to affect both nuclear and cytoplasmic exosome activities. Small RNA-sequencing demonstrated that although some miRNA targets did produce siRNA quantities different from wild type (**Figure 4A**), the *hen2* and *rrp4* mutants exhibited vast differences in their small RNA profiles compared to wild type and *ski2* (**Figure 4B**). In addition to the previously observed siRNA generation from divergent non-coding transcripts (67), many different classes of RNA, in particular mRNA, gave rise to small RNAs (**Figure 4B**). These ectopic small RNAs showed a size distribution typical of siRNAs resulting from DICER-LIKE cleavages with peaks at 21nt and 24 nt (**Figure 4C**), indicating that they are *bona fide* siRNAs and not simply random degradation fragments. Because of the many mRNAs present in the siRNA-producing set, we also tested whether the Zn-finger protein SOP1, related to the key component of the mammalian nuclear Poly(A)-directed exosome targeting complex, ZCCHC1 (68), showed similar deregulated siRNA production. Unlike *hen2* and *rrp4* mutants, however, *sop1-5* did not exhibit substantial ectopic siRNA production (**Supplementary Figure 4**), precluding SOP1-dependent nuclear (pre)-mRNA decay as the major pathway required to limit massive mRNA-derived siRNAs observed in *hen2* and *rrp4*. We also included a mutant in the nuclear exosome subunit RRP45A in our analysis, but did not observe ectopic siRNAs, in this case probably because of unnatural replacement by the more highly expressed RRP45B in the absence of RRP45A protein (**Supplementary Figure 4**).

**Figure 4.**
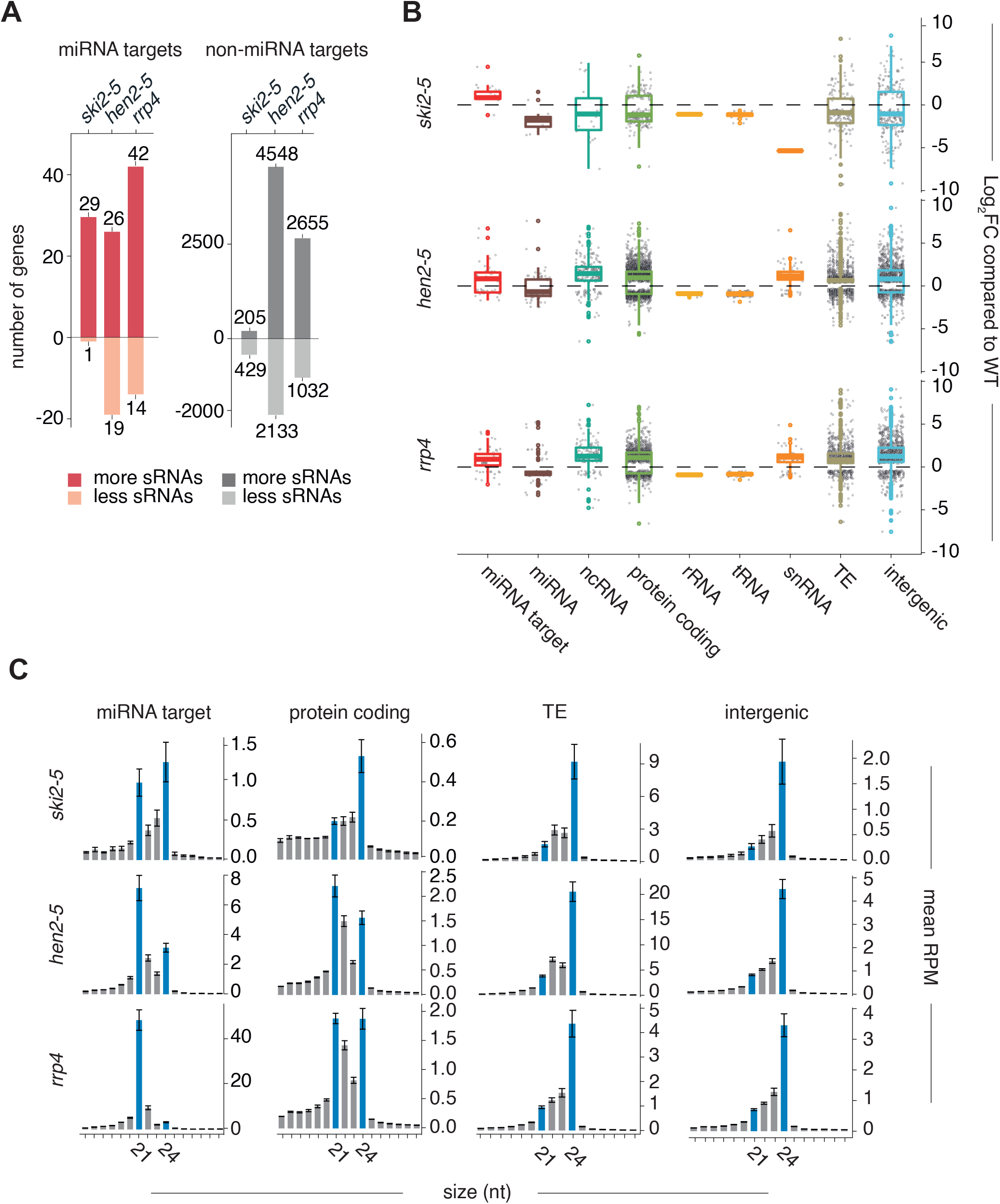
HEN2 and RRP4 have effects on siRNA production from RNAs beyond miRNA targets. (**A**) Bar plots depicting number of known miRNA targets (red bars) or number of non-miRNA targets (grey bars) which produce either more or less secondary siRNAs in *ski2-5, hen2-5* or *rrp4* compared to Col-0 WT (Wald test, P < 0.05). The enrichment of miRNA targets in genes producing more siRNAs is significant in *ski2-5* and *rrp4* (Fisher-test for *ski2-5*: P <2.2 × 10^−16^, Fisher-test for *rrp4*: P=1.7 × 10^−4^). The proportion of miRNA targets found in genes with lower levels of siRNAs in the *ski2-5* and *rrp4* mutants compared to WT is not significant (Fisher-test for *ski2-5*: P = 1.00, Fisher-test for *rrp4:* P= 0.07551). The enrichment of miRNA targets in genes producing both more and less siRNAs in *hen2-5* compared to WT is not significant (Fisher-test for miRNA target enrichment in genes with more siRNAs: P = 0.12, Fisher-test for miRNA target enrichment in genes with less siRNAs: P = 0.62). (**B**) Boxplot of all genes with significantly differentially expressed siRNAs mapped to them (padj. < 0.05). The y-axis shows the log2 of the fold change of read densities (RPM(mutant)/RPM(WT)) of siRNAs mapped to genes. Genes are split into different types according to Araport11 annotations (GFF3 file custom modified to include annotations of intergenic regions). (**C**) Bar plots showing the size distribution of siRNAs mapped to miRNA targets, protein coding genes, transposable elements (TE) and intergenic regions in the different mutants indicated. Error bars show the standard error of the mean RPM in biological triplicates. Small RNA libraries prepared from biological triplicates were used for all analyses presented in this figure. Note that the experiment reported here is fully independent from the one reported in Figure 2 (biological material, number of replicates, library construction), explaining the different number of genes producing siRNAs in *ski2-5* reported in Figure 2 and this figure.

### *rrp4* has a more profound effect on limitation of miRNA-induced secondary siRNAs than either *ski2* or *hen2*

The fact that a high number of mRNAs gives rise to secondary siRNA production in *hen2* and *rrp4* mutants might mean that miRNA targets are included in the set of ectopic siRNA producers by coincidence. If so, the production of siRNAs from these targets would not allow inferences on the possible implication of HEN2 and RRP4 in limitation of miRNA-induced siRNA production. We therefore first inspected the pattern of accumulation of siRNAs mapping to miRNA targets in *hen2* and *rrp4*. Metaplots of siRNA read densities centered on miRNA-guided cleavage sites showed an asymmetry with higher read counts 5’ to cleavage sites and lower read counts 3’ to cleavage sites (**Figure 5A**). This pattern is indicative of a miRNA-RISC-triggered process and is consistent with the fact that most miRNA-triggered secondary siRNAs map 5’ to the cleavage site, perhaps as a consequence of asymmetry in base pairing strength between the 5’ (seed) and 3’ halves of miRNAs to their targets (36). The analysis of siRNAs mapping to miRNA targets in *hen2* and *rrp4* also showed that while siRNA abundances on individual targets tended to be highest close to the miRNA target site, they covered larger parts of target transcripts compared to what is observed in *ski2, pel1* and *rrp45b*, and were generally not restricted to either 5’- or 3’-cleavage fragments (**Figure 5A-C**). We next compared the identities of miRNA target mRNAs giving rise to secondary siRNAs in *ski2, hen2* and *rrp4*. These analyses showed that nearly perfectly reciprocal sets of miRNA targets gave rise to secondary siRNAs in *ski2* and *hen2*, and that the union of those two sets largely matched the set of miRNA targets giving rise to siRNAs in *rrp4* (**Figure 5D, E;; Supplemental Figure 5A,B**). We did not include *hen2/ski2* double mutants in this analysis, because their strong developmental phenotype (**Figure 5F,G**) made it impossible to produce comparable inflorescence tissues for siRNA analysis. A tempting and straightforward interpretation of the observation of reciprocity in miRNA-target/mRNA pairs giving rise to secondary siRNAs between *ski2* and *hen2* mutants is that SKI2 and HEN2 perform biochemically similar functions in limiting miRNA-induced secondary siRNA, with SKI2 acting in the cytoplasm (69), and HEN2 acting in the nucleoplasm (55). Clearly, this interpretation would mean that some miRNA-mRNA targeting events occur preferentially in the cytoplasm while others occur preferentially in the nucleus. If so, the tendency of individual miRNA-targets to produce siRNAs mapping both 5’ and 3’ to miRNA target sites in *hen2* mutants may be explained by a higher tendency of nuclear RDR6 to engage in amplification, including 3’-spreading initiated by a subset of secondary siRNAs. More trivial explanations than nuclear-cytoplasmic partitioning of miRNA-mRNA targeting events are also possible, however. For example, inactivation of HEN2/RRP4 may allow nuclear escape of defective mRNA species and hence provide a pool of mRNA particularly sensitive to RdRP recruitment upon RISC targeting in the cytoplasm.

**Figure 5.**
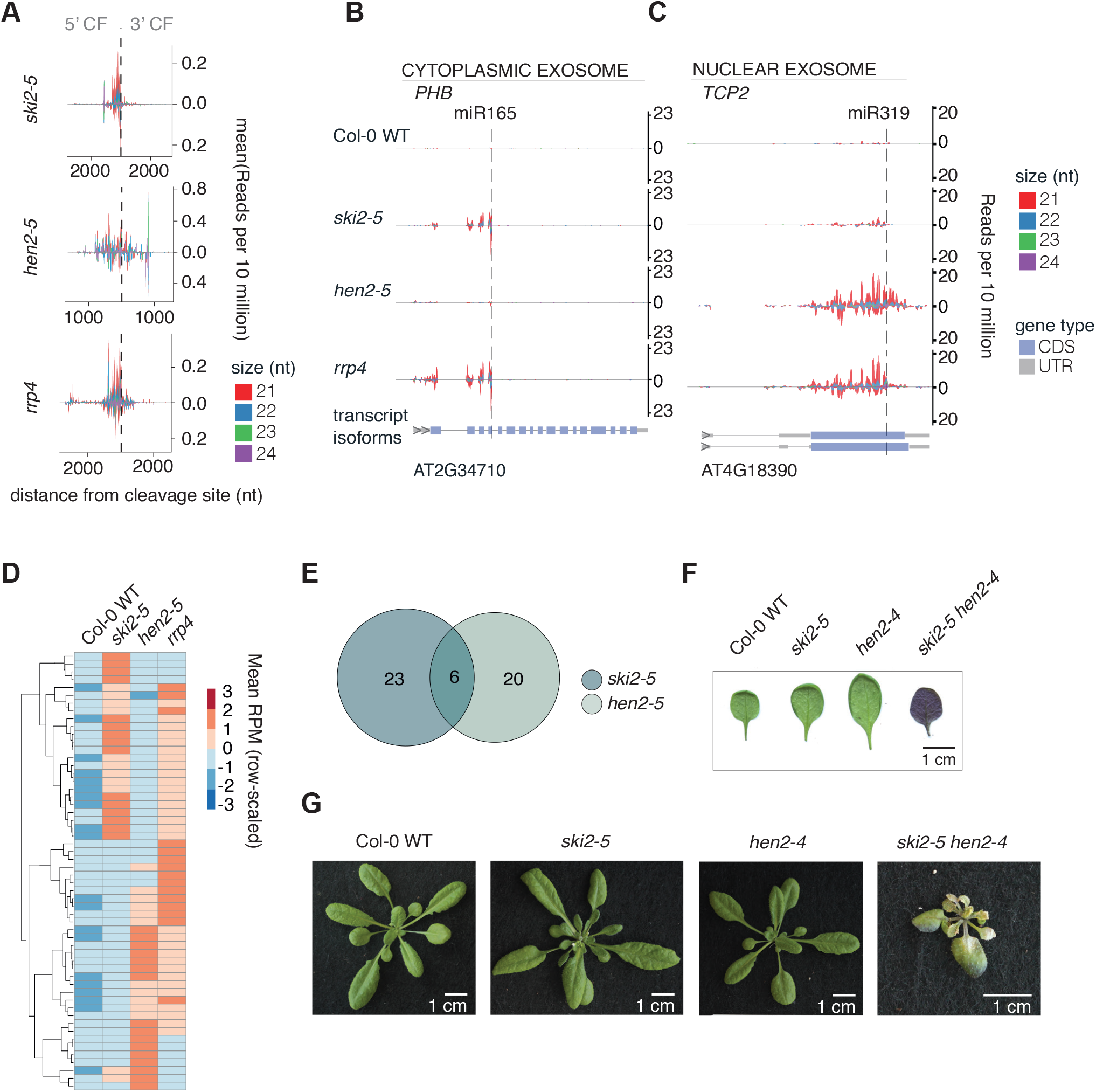
Distinct patterns of miRNA-induced secondary siRNA production in cytoplasmic and nuclear exosome mutants. **(A)** Metaplot of siRNA read densities (RP10M) along miRNA target transcripts with significantly higher siRNA production in mutants than in wild type. Position 0 is defined by miRNA-guided cleavage sites. **(B)** An example of ectopic secondary siRNA accumulation on a known miRNA target, *PHB*, in *ski2-5* and *rrp4* mutants, but not in *hen2-5*. **(C)** An example of ectopic secondary siRNA accumulation from a known miRNA target, *TCP2*, in *hen2-5* and *rrp4* mutants, but not in *ski2-5*. Plots are organized as in Figure 2C and 3B. **(D)** Heatmap of known miRNA targets with significantly more siRNAs produced in *ski2-5, hen2-5* and *rrp4* compared to WT (padj. < 0.05). In the heatmap, the z-score of the mean RPM of siRNAs mapped to each miRNA target in WT, *ski2-5, hen2-5* and *rrp4* are used. The heatmap is clustered by targets. Small RNA libraries prepared from biological triplicates were used for all analyses presented in this figure. (**E**) Euler diagrams showing overlap in siRNA-producing miRNA targets between *ski2-5* and *hen2-5* mutants. (**F**) Abaxial side of detached first true leaves of Col-0, *ski2-5, hen2-4* and *hen2-4/ski2-5*. (**G**) Rosette phenotypes of Col-0, *ski2-5, hen2-4* and *hen2-4/ski2-5* mutants. The plants were photographed after 25 days of growth.

### Illicit secondary siRNAs are not phased

We next analyzed the miRNA-induced siRNA populations in *ski2, hen2* and *rrp4* for phasing relative to the cleavage site, as this may reveal important insight into their mechanism of generation. A phased pattern of accumulation implies that secondary siRNAs resulting from the initial miRNA-dependent recruitment of RDR6 do not trigger further amplification, because small RNA-guided cleavage by AGO1 occurs opposite of nucleotides 10-11 of the guide RNA, thus leading to a 10-nt phase shift if re-amplification occurs (**Figure 6A**). In contrast, multiple scenarios can explain lack of phasing, including reiterative amplification initially starting from a well-defined point, and single-round amplifications starting from template RNAs not perfectly in phase. We detected no phasing of the illicit siRNA populations mapping to PHB and TCP2, in contrast to the phased *TAS1c* siRNAs (**Figure 6B**). At least part of the reason for this result appears to be unaligned 3’-ends of the 5’-cleavage fragments: for PHB, we found that the 3’-most siRNA species, resulting from the first Dicer-catalyzed cleavage event, occurred in at least 6 different phases because of nucleotide tailing and trimming of the PHB 5’-cleavage fragment (**Figure 6C**).

**Figure 6.**
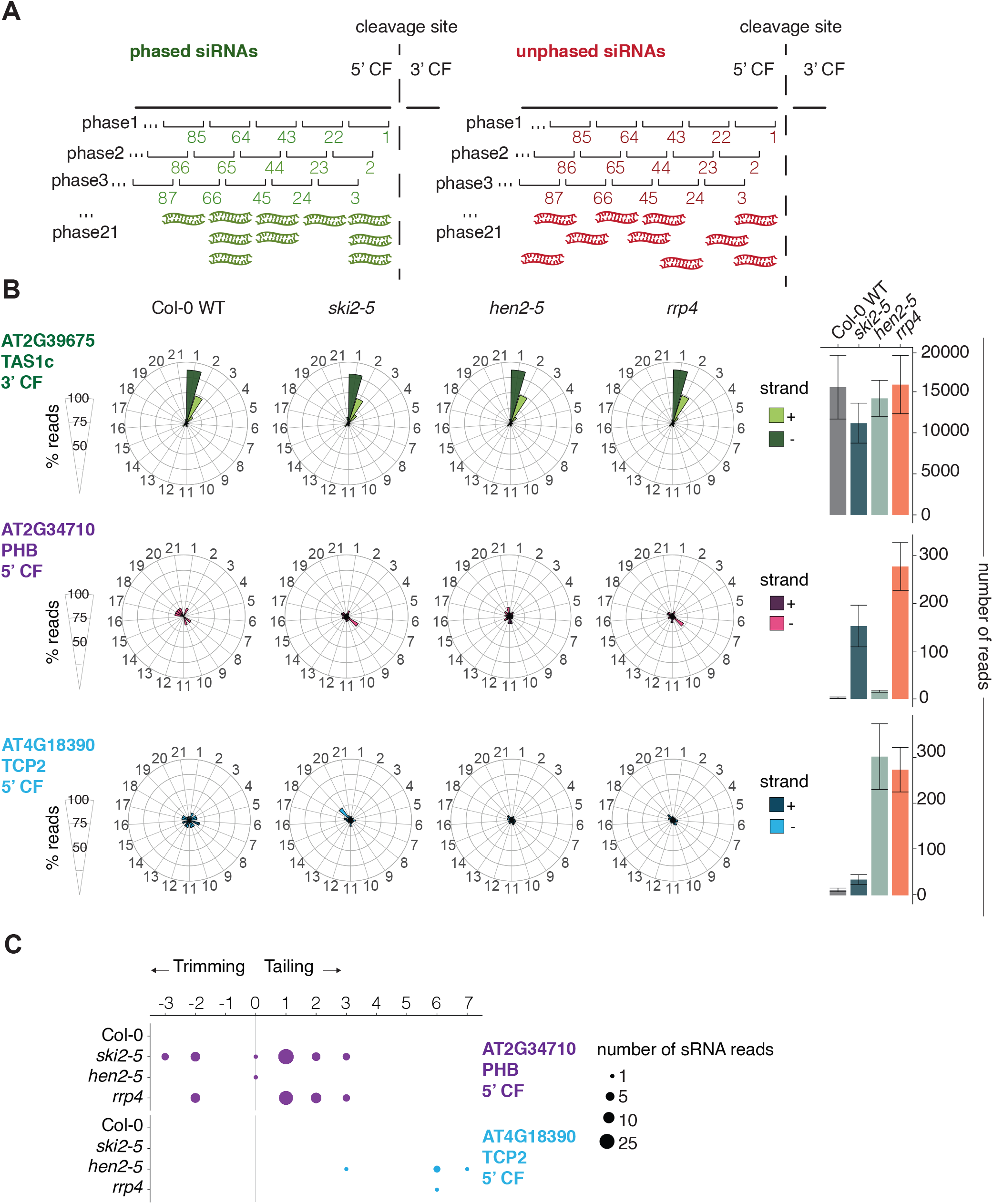
Secondary siRNAs amplified from endogenous miRNA targets in *ski2-5, hen2-4* and *rrp4* mutants are not phased. **(A)** Illustration of phased and unphased siRNA populations, and definitions of the phase registers used for the analysis in B. **(B)** Distribution of siRNAs mapping to *TAS1c, PHB* and *TCP2* in the 21 possible phases in *Col-0, ski2-5, hen2-5* and *rrp4* mutants. *TAS1c* serves as a control known to generate phased siRNA populations. Dark shades, reads mapping to the plus strand;; light shades, reads mapping to the minus strand. Bar plots on the right show the total number of reads underlying the analysis of phase distributions in each genotype. Error bars indicate the standard error of the read count biological triplicates. **(C)** Dot plot representation of heterogenic 5’ ends of the small RNA reads which map to the extremity of the *PHB* and *TCP2* 5’ CF in WT, *ski2-5, hen2-5* and *rrp4*. The circle size represents the number of sRNA reads with a certain number of tailed/trimmed nucleotides found to map to the 5’ CF extremity. Small RNA libraries prepared from biological triplicates were used for all analyses presented in this figure.

### Illicit siRNA production correlates with miRNA abundance, not cleavage fragment abundance

This and our previous study (36) show that occurrence of stable 5’-cleavage fragments does not correlate with production of secondary siRNAs from miRNA targets in *ski2, ski3* and *rrp45b* mutants. For example, *MYB33* (miR159) and *CSD2* (miR398) show stable 5’-cleavage fragments in *ski2* mutants, but no siRNA production, while *AGO1* (miR168) shows a stable 5’-cleavage fragment, but increased siRNA abundances mapping to the 3’-cleavage fragment. With the extended set of miRNA targets found to produce secondary siRNAs in *rrp4* mutants, we therefore asked if other properties of miRNA/target pairs could be identified that correlate with the tendency to initiate secondary siRNAs. We found that the miRNA expression level closely reflects its ability to trigger secondary siRNAs in *rrp4* mutants (**Figure 7A, B**). Above an expression threshold of roughly 250 RPM in *rrp4*, all miRNAs triggered secondary siRNAs, while very few miRNAs below this threshold did so (**Figure 7A**). Similarly, all mRNAs giving rise to log2-fold changes in siRNA abundance higher than 2 were targets of the most highly expressed miRNAs (**Figure 7B**).

**Figure 7.**
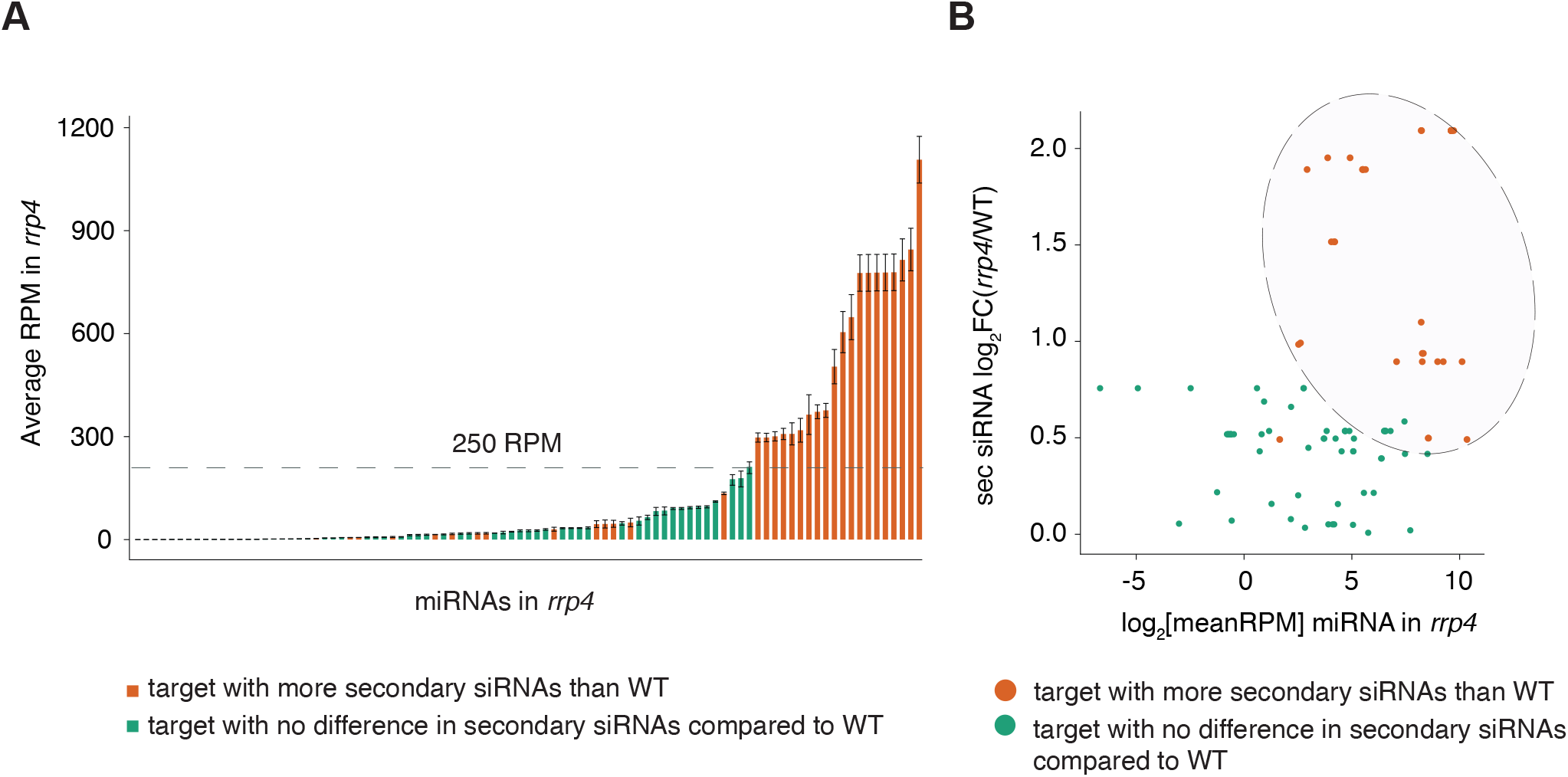
Correlation between miRNA expression level and miRNA-induced secondary siRNA production. **(A)** Bar plot showing expression level (RPM) of miRNAs in *rrp4*. The mean RPM is plotted on the y-axis for each miRNA gene on the x-axis. Error bars show the standard error of the mean RPM of biological *rrp4* triplicates. Red bars, miRNA genes with a significantly higher number of siRNAs in *rrp4* compared to wild type;; green bars, miRNA genes with no significant difference in siRNAs in *rrp4* compared to wild type. The horizontal dashed line indicates an approximate “threshold” of miRNA expression level above which secondary siRNA amplification always is observed. **(B)** Scatterplot showing the relation between miRNA expression level (x-axis) and the fold change value of siRNAs mapped to a target of the miRNA in *rrp4* (y-axis). Red and green colors are employed as in A. The pink circle highlights the tendency of more highly expressed miRNAs to trigger higher levels of secondary siRNAs. Small RNA libraries prepared from biological triplicates were used for all analyses presented in this figure.

## DISCUSSION

### bExosomal degradation of the 5’-cleavage fragment as an accelerator of RISC dissociation

We previously suggested that miRNA-triggered secondary siRNA production involves RISC itself at the target RNA as the key initiating element and termed this model for siRNA amplification the “RISC-trigger model” (36). In contrast, we refer to models that propose initiation of secondary siRNA production through target RNA cleavage fragments by virtue of their aberrant properties as the “aberrant RNA model”. The RISC-trigger model proposes as its central feature that the dwell time of RISC on target RNA is decisive for recruitment of the machinery required for secondary siRNA formation, such that long dwell times favour RdRP recruitment, while rapid RISC dissociation leads to target regulation in the absence of secondary siRNA formation. Evidence consistent with the RISC-trigger, but incompatible with the aberrant RNA model, includes examples of miRNA targets with stable cleavage 5’-cleavage fragments in *ski2* mutants that either fail to produce secondary siRNAs or produce secondary siRNAs mapping to the 3’-cleavage fragment. More direct support for the RISC-trigger model comes from the observation that miR173 triggers secondary siRNAs from *TAS1* and *TAS2* precursors in the complete absence of slicer activity of AGO1 such that no cleavage fragments are generated (60). Likewise, AGO7-miR390 is capable of triggering *TAS3* secondary siRNAs from uncleavable miR390 sites in *TAS3* precursor RNAs (70). In the following, we offer an interpretation of the results described here within the framework of the RISC-trigger model. We stress that despite some attractive features, this model cannot at present be viewed as fully supported by evidence, and we take care to emphasize predictions that arise specifically from interpretations of the present results on the basis of the RISC-trigger model.

The fact that inactivation of the *SKI3* and *RRP45B* genes lead to illicit miRNA-triggered siRNA production very similar to what is observed in *ski2* mutants strongly suggests that there is no previously unrecognized biochemical activity of SKI2 specifically linked to limitation of secondary siRNA production: this effect of SKI2 must also be explained by facilitating exosome function, almost certainly the degradation of 5’-cleavage fragments. But how does failure to degrade 5’-cleavage fragments lead to ectopic siRNA generation 3’ to the cleavage site, as in the example of *AGO1* mRNA? We suggest that SKI2/3/8-exosome-mediated degradation of 5’-cleavage fragments happens *before*, not after, RISC has dissociated from its cleaved target RNA: dynamic “breathing” of base pairs between the 5’-cleavage fragment and the RISC-bound miRNA may allow interaction with SKI2/3/8 while RISC and cleavage fragments are still held together. In this way, exosomal decay of 5’-cleavage fragments would accelerate dissociation of RISC from cleaved target RNAs, and hence maintain dwell times short enough to avoid recruitment of the machinery required for secondary siRNA production. Clearly, such a mechanism predicts physical proximity between SKI2-3-8/exosome and RISC, a property that has not yet been observed in plants, but for which there is precedent in other systems. For example, the *Neurospora crassa* Argonaute protein QDE-2 participates in biogenesis of miRNA-like small RNAs by association with longer precursors such that exosome-mediated trimming of QDE-2-bound small RNA precursors results in mature RISC containing a QDE-2-bound small RNA (71). We also note that the correlation between miRNA levels, rather than cleavage fragment levels, and ability to induce secondary siRNAs in *rrp4* mutants is well explained by the RISC-trigger model. If the absence of SKI2/3/8-exosome activity shifts the distribution of dwell times such that a fraction of RISC-target mRNA encounters is long-lived enough to trigger secondary siRNA formation, the resulting quantity of secondary siRNAs becomes proportional to the number of RISC-target mRNA encounters, and hence to the product of miRNA and target mRNA concentrations. Finally, we note that the RISC-trigger model may provide a simple explanation for the tendency of 22-nt miRNAs to induce siRNA amplification, even in the presence of SKI2-3-8/exosome function: the additional base pair between the target 5’-cleavage fragment and the 3’-end of the miRNA may prolong the average dwell time of RISC sufficiently that at least some RISC:target mRNA encounters lead to recruitment of the siRNA amplification machinery. This interpretation would also explain the recent observation that a 1-nt insertion mutant in miR398b to make it 22-nt long does not induce secondary siRNAs (72). In the RISC-trigger model, the mutant 22-nt miR398b does not induce siRNA amplification, because the additional nucleotide in the miRNA does not result in formation of additional target base pairs, and hence does not lead to long-lived miR398b-RISC:target interactions.

### Distinct roles of PEL1 and SKI2 in limiting secondary siRNA production

PEL1, SKI2-3-8 and the cytoplasmic exosome have a clear role in degradation of RISC 5’-cleavage fragments and other NSD substrates (36,50). Yet, as discussed at length above, the stable 5’-cleavage fragments are not direct RdRP substrates, suggesting that the requirement for PEL1 for avoidance of miRNA-triggered siRNAs may not be related to degradation of 5’-cleavage fragments, and, by consequence, that inactivation of PEL1 and SKI2-3-8-exosome leads to miRNA-triggered siRNA production for different reasons. Consistent with this idea, the set of miRNA targets that produces siRNAs in *pel1* and *ski2* mutants is not the same, even if there is some overlap. How could loss of PEL1 function stimulate miRNA-induced siRNA production? And in particular, what is the explanation for enhanced siRNA production in *pel1* mutants not only of siRNAs mapping to 5’-cleavage fragments, but also to 3’-cleavage fragments as shown by the miR472-*RSG2* example? Several observations indicate the importance of ribosome association for miRNA-triggered siRNA production from *TAS* precursors. Ribosome profiling experiments demonstrate ribosome association with *TAS* precursors (73,74), and loss of the exportin-like protein SDE5 required for all tasiRNA production causes loss of *TAS* precursor transcripts from ribosome-associated fractions (75). Furthermore, detailed biochemical analyses reveal the importance of ribosome stalling in proximity to the 5’ miR390 site in *TAS3* precursors (74), and ribosome stalling and collision at rare codons in transposable element mRNAs correlates with their production of 21-nt siRNAs (53) (observed in mutants defective in DNA methylation, and, therefore, often referred to as epigenetically activated siRNAs, or easiRNAs (25)). It appears, therefore, that stalled ribosomes, in particular in combination with RISC, act as a trigger of secondary siRNA production, perhaps by extending the time of RISC association with target mRNA by AGO interaction. Indeed, AGO proteins, including Arabidopsis AGO1, associate with polyribosomes and co-purify with ribosomal proteins (76-78). Thus, the subunit dissociation activity on stalled ribosomes of Dom34/Pelota and Hbs1 (51,79) could directly limit secondary siRNA formation from miRNA-target mRNAs associated with RISC. This proposition is consistent with all of the properties of enhanced miRNA-induced siRNA production in *pel1* mutants that we observe here, in particular the distinct set of miRNA targets affected compared with *ski2* mutants, and enhanced production of siRNAs mapping to 3’-cleavage fragments in some instances. We note that this model requires plant PEL1 to be able to cause splitting of subunits from ribosomes stalled at internal sites in mRNAs. It is at present unclear to what extent internal ribosome stalls are resolved by Pelota:Hbs1 (42,44,51,79), particularly because Pelota binding to stalled ribosomes is favoured by A-sites free of mRNA (51,80). Nonetheless, cryo-EM structures of Pelota bound to stalled ribosomes with mRNA sequence downstream of the P-site indicated mRNA displacement from the channel upon Pelota binding, thus strongly suggesting that an empty A-site is not an absolute requirement for Pelota binding (81). In addition, as noted by D’Orazio and Green (42), the observation that loss of the mouse Hbs1 homologue GTPBP2 leads to neurodegeneration in a mouse strain carrying an inactivating mutation in a brain-specific Arg-tRNA provides an example of requirement of resolution of internal ribosome stalls *in vivo* by Pelota:Hbs1 (82). Thus, although a direct implication of stalled ribosome eviction by plant PEL1 in limitation of secondary siRNA production is not uniformly supported by biochemical analysis of Pelota/Dom34 activity on stalled ribosomes *in vitro*, this proposition is consistent with other analyses of molecular properties of Hbs1:Pelota and their function *in vivo*.

### Possible reasons for ARGONAUTE specialization in secondary siRNA formation

The most highly conserved example of miRNA-induced secondary siRNA production in plants is the formation of AUXIN RESPONSE FACTOR (ARF)-targeting *TAS3* siRNAs by miR390. These tasiRNAs are fundamental for leaf and flower development, as documented by consequences of their loss in mutants required specifically for siRNA amplification. These defects include aberrant developmental timing (83), and leaf and flower morphogenesis defects in multiple species (84-89). *TAS3* siRNAs are initiated by a conserved AGO7-miR390 RISC (90), not by an AGO1-based RISC, and it has not been clearly established why a specialized AGO protein is required for this process of secondary siRNA formation. Our analysis of illicit secondary siRNAs produced by AGO1-miRNA targets may reveal part of the answer to this mystery: in most angiosperms, *TAS3* siRNAs derive from the 5’-cleavage fragment of the *TAS3* precursor (91), by dicing from its 3’-end following miR390-guided cleavage and conversion into dsRNA. The 5’-cleavage fragment must therefore maintain a well-defined 3’-end to produce functional ARF-targeting siRNAs in phase with the miR390-guided cleavage site. However, AGO1-miR166 induced secondary siRNAs mapping to *PHB*, and AGO1-miR319 induced secondary siRNAs mapping to *TCP2* were completely unphased, at least in part due to tailing and nibbling of the 3’-end of the 5’-cleavage fragment. Strong tailing and trimming activities are associated with AGO1, but less so with AGO7, as suggested by the different effect of mutation of the small RNA methyl transferase HEN1 on AGO1-bound miR173 (and other miRNAs) and AGO7-bound miR390. In *hen1* mutants, AGO1-bound miRNAs decrease dramatically in abundance and accumulate as a distribution of 17-27 nt species, while miR390 has reduced abundance, but largely maintains a 21-nt size (92). Thus, employment of a specialized AGO protein, AGO7, may help maintain the crucial phased register of *TAS3* siRNAs. We note, however, that this cannot be the only specialization of AGO7 relevant for *TAS3* siRNA biogenesis, as previous analyses also pointed to the requirement of AGO7-specific activities at the non-cleavable miR390 site in *TAS3* precursors (90).

## AVAILABILITY

The sequencing data generated in this publication have been deposited in NCBI’s Gene Expression Omnibus (93) (https://www.ncbi.nlm.nih.gov/geo/query/acc.cgi?acc=GSE173426)

R code is available in the GitHub repository https://github.com/MariaLouisaVigh/ExosomePelota.

## ACCESSION NUMBERS

Small RNA sequencing data are accessible through GEO Series accession number GSE173426.

## SUPPLEMENTARY MATERIAL

This article contains supplementary data:

Supplementary Figure 1: **Validation of small RNA sequencing data prior to DESeq analyses**.

Supplementary Figure 2: **miRNA expression levels in mutants used in this study compared to Col-0 WT**.

Supplementary Figure 3: **miRNA-triggered siRNA accumulation in *ski2-5* and *pel1* mutants**

Supplementary Figure 4: **siRNA accumulation in *rrp45a* and *sop1-5* mutants**.

Supplementary Figure 5: **siRNA production from miRNA targets in *ski2-5, hen2-5* and *rrp4-2***

Supplementary Table 1: **List of oligonucleotides used in the study**

Supplementary Table 2: **List of genes with significant different levels of sRNAs in *ski2-5, ski3-5* and *rrp45b* (Experiment A)**

Supplementary Table 3: **List of genes with significant different levels of sRNAs in *ski2-5, pel1-1, pel1-2, hen2-5, rrp4, rrp45a* (Experiment B)**

Supplementary Table 4: **List of genes with significant different levels of sRNAs in *ski2-5, hen2-5* and *sop1-5* (Experiment C)**

Supplementary Table 5: **List of known miRNA targets (experimentally verified)**

## ACKNOWLEDGMENTS

We thank the greenhouse teams of Theo Bølsterli and René Hvidberg for plant care. Lena Bjørn Johansson is thanked for help with the northern blot analyses, and Andrea Barghetti is thanked for help with data analysis. Emilie Oksbjerg is thanked for constructive scientific discussion, and Diego López-Márquez is thanked for critical reading of the manuscript. Ljerka Kunst, Dominique Gagliardi and Kian Hématy are thanked for providing seeds of *cer7-3, cer7-3/rdr6-12* (L.K.), *hen2-4* (D.G.), *hen2-5, sop1-5* and *rrp4-2/sop2* (K.H).

## REFERENCES

1. Fang, X. and Qi, Y. (2016) RNAi in Plants: An Argonaute-Centered View. Plant Cell, 28, 272–285.

2. Meister, G. (2013) Argonaute proteins: functional insights and emerging roles. Nat Rev Genet, 14, 447–459.

3. Liu, Y., Teng, C., Xia, R. and Meyers, B.C. (2020) PhasiRNAs in Plants: Their Biogenesis, Genic Sources, and Roles in Stress Responses, Development, and Reproduction. Plant Cell, 32, 3059–3080.

4. Svoboda, P. (2020) Key Mechanistic Principles and Considerations Concerning RNA Interference. Front Plant Sci, 11, 1237.

5. Pak, J. and Fire, A. (2007) Distinct populations of primary and secondary effectors during RNAi in C. elegans. Science, 315, 241–244.

6. Moissiard, G., Parizotto, E.A., Himber, C. and Voinnet, O. (2007) Transitivity in Arabidopsis can be primed, requires the redundant action of the antiviral Dicer-like 4 and Dicer-like 2, and is compromised by viral-encoded suppressor proteins. RNA, 13, 1268–1278.

7. Yoshikawa, M., Peragine, A., Park, M.Y. and Poethig, R.S. (2005) A pathway for the biogenesis of trans-acting siRNAs in Arabidopsis. Genes Dev, 19, 2164–2175.

8. Xie, Z., Allen, E., Wilken, A. and Carrington, J.C. (2005) DICER-LIKE 4 functions in transacting small interfering RNA biogenesis and vegetative phase change in Arabidopsis thaliana. Proc Natl Acad Sci U S A, 102, 12984–12989.

9. Parizotto, E.A., Dunoyer, P., Rahm, N., Himber, C. and Voinnet, O. (2004) In vivo investigation of the transcription, processing, endonucleolytic activity, and functional relevance of the spatial distribution of a plant miRNA. Genes & development, 18, 2237–2242.

10. Carlsbecker, A., Lee, J.Y., Roberts, C.J., Dettmer, J., Lehesranta, S., Zhou, J., Lindgren, O., Moreno-Risueno, M.A., Vaten, A., Thitamadee, S. et al. (2010) Cell signalling by microRNA165/6 directs gene dose-dependent root cell fate. Nature, 465, 316–321.

11. Brosnan, C.A., Sarazin, A., Lim, P., Bologna, N.G., Hirsch-Hoffmann, M. and Voinnet, O. (2019) Genome-scale, single-cell-type resolution of microRNA activities within a whole plant organ. EMBO J, 38, e100754.

12. de Felippes, F.F., Ott, F. and Weigel, D. (2011) Comparative analysis of non-autonomous effects of tasiRNAs and miRNAs in Arabidopsis thaliana. Nucleic Acids Res, 39, 2880–2889.

13. Tsikou, D., Yan, Z., Holt, D.B., Abel, N.B., Reid, D.E., Madsen, L.H., Bhasin, H., Sexauer, M., Stougaard, J. and Markmann, K. (2018) Systemic control of legume susceptibility to rhizobial infection by a mobile microRNA. Science, 362, 233–236.

14. Pant, B.D., Buhtz, A., Kehr, J. and Scheible, W.R. (2008) MicroRNA399 is a long-distance signal for the regulation of plant phosphate homeostasis. Plant J, 53, 731–738.

15. Allen, E., Xie, Z., Gustafson, A.M. and Carrington, J.C. (2005) microRNA-directed phasing during trans-acting siRNA biogenesis in plants. Cell, 121, 207–221.

16. Klesen, S., Hill, K. and Timmermans, M.C.P. (2020) Small RNAs as plant morphogens. Curr Top Dev Biol, 137, 455–480.

17. Skopelitis, D.S., Benkovics, A.H., Husbands, A.Y. and Timmermans, M.C.P. (2017) Boundary Formation through a Direct Threshold-Based Readout of Mobile Small RNA Gradients. Dev Cell, 43, 265–273 e266.

18. Cai, Q., Qiao, L., Wang, M., He, B., Lin, F.M., Palmquist, J., Huang, S.D. and Jin, H. (2018) Plants send small RNAs in extracellular vesicles to fungal pathogen to silence virulence genes. Science, 360, 1126–1129.

19. Hou, Y., Zhai, Y., Feng, L., Karimi, H.Z., Rutter, B.D., Zeng, L., Choi, D.S., Zhang, B., Gu, W., Chen, X. et al. (2019) A Phytophthora Effector Suppresses Trans-Kingdom RNAi to Promote Disease Susceptibility. Cell Host Microbe, 25, 153–165 e155.

20. Dalmay, T., Hamilton, A., Rudd, S., Angell, S. and Baulcombe, D.C. (2000) An RNA-dependent RNA polymerase gene in Arabidopsis is required for posttranscriptional gene silencing mediated by a transgene but not by a virus. Cell, 101, 543–553.

21. Mourrain, P., Beclin, C., Elmayan, T., Feuerbach, F., Godon, C., Morel, J.B., Jouette, D., Lacombe, A.M., Nikic, S., Picault, N. et al. (2000) Arabidopsis SGS2 and SGS3 genes are required for posttranscriptional gene silencing and natural virus resistance. Cell, 101, 533–542.

22. Dalmay, T., Horsefield, R., Braunstein, T.H. and Baulcombe, D.C. (2001) SDE3 encodes an RNA helicase required for post-transcriptional gene silencing in Arabidopsis. EMBO J, 20, 2069–2078.

23. Garcia-Ruiz, H., Takeda, A., Chapman, E.J., Sullivan, C.M., Fahlgren, N., Brempelis, K.J. and Carrington, J.C. (2010) Arabidopsis RNA-dependent RNA polymerases and dicer-like proteins in antiviral defense and small interfering RNA biogenesis during Turnip Mosaic Virus infection. Plant Cell, 22, 481–496.

24. Mari-Ordonez, A., Marchais, A., Etcheverry, M., Martin, A., Colot, V. and Voinnet, O. (2013) Reconstructing de novo silencing of an active plant retrotransposon. Nat Genet, 45, 1029–1039.

25. Creasey, K.M., Zhai, J., Borges, F., Van Ex, F., Regulski, M., Meyers, B.C. and Martienssen, R.A. (2014) miRNAs trigger widespread epigenetically activated siRNAs from transposons in Arabidopsis. Nature, 508, 411–415.

26. Baeg, K., Iwakawa, H.O. and Tomari, Y. (2017) The poly(A) tail blocks RDR6 from converting self mRNAs into substrates for gene silencing. Nat Plants, 3, 17036.

27. Chen, H.M., Chen, L.T., Patel, K., Li, Y.H., Baulcombe, D.C. and Wu, S.H. (2010) 22-Nucleotide RNAs trigger secondary siRNA biogenesis in plants. Proc Natl Acad Sci U S A, 107, 15269–15274.

28. Cuperus, J.T., Carbonell, A., Fahlgren, N., Garcia-Ruiz, H., Burke, R.T., Takeda, A., Sullivan, C.M., Gilbert, S.D., Montgomery, T.A. and Carrington, J.C. (2010) Unique functionality of 22-nt miRNAs in triggering RDR6-dependent siRNA biogenesis from target transcripts in Arabidopsis. Nat Struct Mol Biol, 17, 997–1003.

29. Fei, Q., Yu, Y., Liu, L., Zhang, Y., Baldrich, P., Dai, Q., Chen, X. and Meyers, B.C. (2018) Biogenesis of a 22-nt microRNA in Phaseoleae species by precursor-programmed uridylation. Proc Natl Acad Sci U S A, 115, 8037–8042.

30. Gy, I., Gasciolli, V., Lauressergues, D., Morel, J.B., Gombert, J., Proux, F., Proux, C., Vaucheret, H. and Mallory, A.C. (2007) Arabidopsis FIERY1, XRN2, and XRN3 are endogenous RNA silencing suppressors. Plant Cell, 19, 3451–3461.

31. Moreno, A.B., Martinez de Alba, A.E., Bardou, F., Crespi, M.D., Vaucheret, H., Maizel, A. and Mallory, A.C. (2013) Cytoplasmic and nuclear quality control and turnover of single-stranded RNA modulate post-transcriptional gene silencing in plants. Nucleic Acids Res, 41, 4699–4708.

32. Scheer, H., de Almeida, C., Ferrier, E., Simonnot, Q., Poirier, L., Pflieger, D., Sement, F.M., Koechler, S., Piermaria, C., Krawczyk, P. et al. (2021) The TUTase URT1 connects decapping activators and prevents the accumulation of excessively deadenylated mRNAs to avoid siRNA biogenesis. Nat Commun, 12, 1298.

33. Lange, H., Ndecky, S.Y.A., Gomez-Diaz, C., Pflieger, D., Butel, N., Zumsteg, J., Kuhn, L., Piermaria, C., Chicher, J., Christie, M. et al. (2019) RST1 and RIPR connect the cytosolic RNA exosome to the Ski complex in Arabidopsis. Nat Commun, 10, 3871.

34. Martinez de Alba, A.E., Moreno, A.B., Gabriel, M., Mallory, A.C., Christ, A., Bounon, R., Balzergue, S., Aubourg, S., Gautheret, D., Crespi, M.D. et al. (2015) In plants, decapping prevents RDR6-dependent production of small interfering RNAs from endogenous mRNAs. Nucleic Acids Res, 43, 2902–2913.

35. Li, T., Natran, A., Chen, Y., Vercruysse, J., Wang, K., Gonzalez, N., Dubois, M. and Inze, D. (2019) A genetics screen highlights emerging roles for CPL3, RST1 and URT1 in RNA metabolism and silencing. Nat Plants, 5, 539–550.

36. Branscheid, A., Marchais, A., Schott, G., Lange, H., Gagliardi, D., Andersen, S.U., Voinnet, O. and Brodersen, P. (2015) SKI2 mediates degradation of RISC 5’-cleavage fragments and prevents secondary siRNA production from miRNA targets in Arabidopsis. Nucleic Acids Res, 43, 10975–10988.

37. Anderson, J.S. and Parker, R.P. (1998) The 3’ to 5’ degradation of yeast mRNAs is a general mechanism for mRNA turnover that requires the SKI2 DEVH box protein and 3’ to 5’ exonucleases of the exosome complex. EMBO J, 17, 1497–1506.

38. Brown, J.T., Bai, X. and Johnson, A.W. (2000) The yeast antiviral proteins Ski2p, Ski3p, and Ski8p exist as a complex in vivo. RNA, 6, 449–457.

39. Synowsky, S.A. and Heck, A.J. (2008) The yeast Ski complex is a hetero-tetramer. Protein Sci, 17, 119–125.

40. Halbach, F., Reichelt, P., Rode, M. and Conti, E. (2013) The yeast ski complex: crystal structure and RNA channeling to the exosome complex. Cell, 154, 814–826.

41. Schmid, M. and Jensen, T.H. (2008) The exosome: a multipurpose RNA-decay machine. Trends Biochem Sci, 33, 501–510.

42. D’Orazio, K.N. and Green, R. (2021) Ribosome states signal RNA quality control. Mol Cell, 81, 1372–1383.

43. van Hoof, A., Frischmeyer, P.A., Dietz, H.C. and Parker, R. (2002) Exosome-mediated recognition and degradation of mRNAs lacking a termination codon. Science, 295, 2262–2264.

44. Tsuboi, T., Kuroha, K., Kudo, K., Makino, S., Inoue, E., Kashima, I. and Inada, T. (2012) Dom34:hbs1 plays a general role in quality-control systems by dissociation of a stalled ribosome at the 3’ end of aberrant mRNA. Mol Cell, 46, 518–529.

45. Shoemaker, C.J. and Green, R. (2012) Translation drives mRNA quality control. Nat Struct Mol Biol, 19, 594–601.

46. Doma, M.K. and Parker, R. (2006) Endonucleolytic cleavage of eukaryotic mRNAs with stalls in translation elongation. Nature, 440, 561–564.

47. Jones-Rhoades, M.W. and Bartel, D.P. (2004) Computational identification of plant microRNAs and their targets, including a stress-induced miRNA. Mol Cell, 14, 787–799.

48. Addo-Quaye, C., Eshoo, T.W., Bartel, D.P. and Axtell, M.J. (2008) Endogenous siRNA and miRNA targets identified by sequencing of the Arabidopsis degradome. Curr Biol, 18, 758–762.

49. German, M.A., Pillay, M., Jeong, D.H., Hetawal, A., Luo, S., Janardhanan, P., Kannan, V., Rymarquis, L.A., Nobuta, K., German, R. et al. (2008) Global identification of microRNA-target RNA pairs by parallel analysis of RNA ends. Nat Biotechnol, 26, 941–946.

50. Szadeczky-Kardoss, I., Csorba, T., Auber, A., Schamberger, A., Nyiko, T., Taller, J., Orban, T.I., Burgyan, J. and Silhavy, D. (2018) The nonstop decay and the RNA silencing systems operate cooperatively in plants. Nucleic Acids Res, 46, 4632–4648.

51. Pisareva, V.P., Skabkin, M.A., Hellen, C.U., Pestova, T.V. and Pisarev, A.V. (2011) Dissociation by Pelota, Hbs1 and ABCE1 of mammalian vacant 80S ribosomes and stalled elongation complexes. EMBO J, 30, 1804–1817.

52. Szadeczky-Kardoss, I., Gal, L., Auber, A., Taller, J. and Silhavy, D. (2018) The No-go decay system degrades plant mRNAs that contain a long A-stretch in the coding region. Plant Sci, 275, 19–27.

53. Kim, E.Y., Wang, L., Lei, Z., Li, H., Fan, W. and Cho, J. (2021) Ribosome stalling and SGS3 phase separation prime the epigenetic silencing of transposons. Nat Plants, 7, 303–309.

54. Lange, H., Sement, F.M. and Gagliardi, D. (2011) MTR4, a putative RNA helicase and exosome co-factor, is required for proper rRNA biogenesis and development in Arabidopsis thaliana. Plant J, 68, 51–63.

55. Lange, H., Zuber, H., Sement, F.M., Chicher, J., Kuhn, L., Hammann, P., Brunaud, V., Berard, C., Bouteiller, N., Balzergue, S. et al. (2014) The RNA helicases AtMTR4 and HEN2 target specific subsets of nuclear transcripts for degradation by the nuclear exosome in Arabidopsis thaliana. PLoS Genet, 10, e1004564.

56. Hematy, K., Bellec, Y., Podicheti, R., Bouteiller, N., Anne, P., Morineau, C., Haslam, R.P., Beaudoin, F., Napier, J.A., Mockaitis, K. et al. (2016) The Zinc-Finger Protein SOP1 Is Required for a Subset of the Nuclear Exosome Functions in Arabidopsis. PLoS Genet, 12, e1005817.

57. Kleinboelting, N., Huep, G., Kloetgen, A., Viehoever, P. and Weisshaar, B. (2012) GABI-Kat SimpleSearch: new features of the Arabidopsis thaliana T-DNA mutant database. Nucleic Acids Res, 40, D1211–1215.

58. Hooker, T.S., Lam, P., Zheng, H. and Kunst, L. (2007) A core subunit of the RNA-processing/degrading exosome specifically influences cuticular wax biosynthesis in Arabidopsis. Plant Cell, 19, 904–913.

59. Rosso, M.G., Li, Y., Strizhov, N., Reiss, B., Dekker, K. and Weisshaar, B. (2003) An Arabidopsis thaliana T-DNA mutagenized population (GABI-Kat) for flanking sequence tag-based reverse genetics. Plant Mol Biol, 53, 247–259.

60. Arribas-Hernandez, L., Marchais, A., Poulsen, C., Haase, B., Hauptmann, J., Benes, V., Meister, G. and Brodersen, P. (2016) The Slicer Activity of ARGONAUTE1 Is Required Specifically for the Phasing, Not Production, of Trans-Acting Short Interfering RNAs in Arabidopsis. Plant Cell, 28, 1563–1580.

61. Martin, M. (2011) Cutadapt removes adapter sequences from high-throughput sequencing reads. 2011, 17, 3.

62. Dobin, A., Davis, C.A., Schlesinger, F., Drenkow, J., Zaleski, C., Jha, S., Batut, P., Chaisson, M. and Gingeras, T.R. (2013) STAR: ultrafast universal RNA-seq aligner. Bioinformatics, 29, 15–21.

63. Liao, Y., Smyth, G.K. and Shi, W. (2014) featureCounts: an efficient general purpose program for assigning sequence reads to genomic features. Bioinformatics, 30, 923–930.

64. Love, M.I., Huber, W. and Anders, S. (2014) Moderated estimation of fold change and dispersion for RNA-seq data with DESeq2. Genome Biol, 15, 550.

65. Anders, S. and Huber, W. (2010) Differential expression analysis for sequence count data. Genome Biol, 11, R106.

66. Rainer, J., Gatto, L. and Weichenberger, C.X. (2019) ensembldb: an R package to create and use Ensembl-based annotation resources. Bioinformatics, 35, 3151–3153.

67. Thieffry, A., Vigh, M.L., Bornholdt, J., Ivanov, M., Brodersen, P. and Sandelin, A. (2020) Characterization of Arabidopsis thaliana Promoter Bidirectionality and Antisense RNAs by Inactivation of Nuclear RNA Decay Pathways. Plant Cell, 32, 1845–1867.

68. Meola, N., Domanski, M., Karadoulama, E., Chen, Y., Gentil, C., Pultz, D., Vitting-Seerup, K., Lykke-Andersen, S., Andersen, J.S., Sandelin, A. et al. (2016) Identification of a Nuclear Exosome Decay Pathway for Processed Transcripts. Mol Cell, 64, 520–533.

69. Zhang, X., Zhu, Y., Liu, X., Hong, X., Xu, Y., Zhu, P., Shen, Y., Wu, H., Ji, Y., Wen, X. et al. (2015) Plant biology. Suppression of endogenous gene silencing by bidirectional cytoplasmic RNA decay in Arabidopsis. Science, 348, 120–123.

70. de Felippes, F.F., Marchais, A., Sarazin, A., Oberlin, S. and Voinnet, O. (2017) A single miR390 targeting event is sufficient for triggering TAS3-tasiRNA biogenesis in Arabidopsis. Nucleic Acids Res, 45, 5539–5554.

71. Xue, Z., Yuan, H., Guo, J. and Liu, Y. (2012) Reconstitution of an Argonaute-dependent small RNA biogenesis pathway reveals a handover mechanism involving the RNA exosome and the exonuclease QIP. Mol Cell, 46, 299–310.

72. Bi, H., Fei, Q., Li, R., Liu, B., Xia, R., Char, S.N., Meyers, B.C. and Yang, B. (2020) Disruption of miRNA sequences by TALENs and CRISPR/Cas9 induces varied lengths of miRNA production. Plant Biotechnol J, 18, 1526–1536.

73. Bazin, J., Baerenfaller, K., Gosai, S.J., Gregory, B.D., Crespi, M. and Bailey-Serres, J. (2017) Global analysis of ribosome-associated noncoding RNAs unveils new modes of translational regulation. Proc Natl Acad Sci U S A, 114, E10018–E10027.

74. Iwakawa, H.-o., Lam, A.Y.W., Mine, A., Fujita, T., Kiyokawa, K., Yoshikawa, M., Takeda, A., Iwasaki, S. and Tomari, Y. (2020) Ribosome stalling caused by the Argonaute-miRNA-SGS3 complex regulates production of secondary siRNA biogenesis in plants. bioRxiv, 2020.2009.2010.288902.

75. Yoshikawa, M., Iki, T., Numa, H., Miyashita, K., Meshi, T. and Ishikawa, M. (2016) A Short Open Reading Frame Encompassing the MicroRNA173 Target Site Plays a Role in trans-Acting Small Interfering RNA Biogenesis. Plant Physiol, 171, 359–368.

76. Lanet, E., Delannoy, E., Sormani, R., Floris, M., Brodersen, P., Crete, P., Voinnet, O. and Robaglia, C. (2009) Biochemical evidence for translational repression by Arabidopsis microRNAs. Plant Cell, 21, 1762–1768.

77. Frohn, A., Eberl, H.C., Stohr, J., Glasmacher, E., Rudel, S., Heissmeyer, V., Mann, M. and Meister, G. (2012) Dicer-dependent and -independent Argonaute2 protein interaction networks in mammalian cells. Mol Cell Proteomics, 11, 1442–1456.

78. Ma, X., Ibrahim, F., Kim, E.J., Shaver, S., Becker, J., Razvi, F., Cerny, R.L. and Cerutti, H. (2020) An ortholog of the Vasa intronic gene is required for small RNA-mediated translation repression in Chlamydomonas reinhardtii. Proc Natl Acad Sci U S A, 117, 761–770.

79. Shoemaker, C.J., Eyler, D.E. and Green, R. (2010) Dom34:Hbs1 promotes subunit dissociation and peptidyl-tRNA drop-off to initiate no-go decay. Science, 330, 369–372.

80. Becker, T., Armache, J.P., Jarasch, A., Anger, A.M., Villa, E., Sieber, H., Motaal, B.A., Mielke, T., Berninghausen, O. and Beckmann, R. (2011) Structure of the no-go mRNA decay complex Dom34-Hbs1 bound to a stalled 80S ribosome. Nat Struct Mol Biol, 18, 715–720.

81. Shao, S., Murray, J., Brown, A., Taunton, J., Ramakrishnan, V. and Hegde, R.S. (2016) Decoding Mammalian Ribosome-mRNA States by Translational GTPase Complexes. Cell, 167, 1229–1240 e1215.

82. Ishimura, R., Nagy, G., Dotu, I., Zhou, H., Yang, X.L., Schimmel, P., Senju, S., Nishimura, Y., Chuang, J.H. and Ackerman, S.L. (2014) RNA function. Ribosome stalling induced by mutation of a CNS-specific tRNA causes neurodegeneration. Science, 345, 455–459.

83. Hunter, C., Willmann, M.R., Wu, G., Yoshikawa, M., de la Luz Gutierrez-Nava, M. and Poethig, S.R. (2006) Trans-acting siRNA-mediated repression of ETTIN and ARF4 regulates heteroblasty in Arabidopsis. Development, 133, 2973–2981.

84. Garcia, D., Collier, S.A., Byrne, M.E. and Martienssen, R.A. (2006) Specification of leaf polarity in Arabidopsis via the trans-acting siRNA pathway. Curr Biol, 16, 933–938.

85. Fahlgren, N., Montgomery, T.A., Howell, M.D., Allen, E., Dvorak, S.K., Alexander, A.L. and Carrington, J.C. (2006) Regulation of AUXIN RESPONSE FACTOR3 by TAS3 ta-siRNA affects developmental timing and patterning in Arabidopsis. Curr Biol, 16, 939–944.

86. Adenot, X., Elmayan, T., Lauressergues, D., Boutet, S., Bouche, N., Gasciolli, V. and Vaucheret, H. (2006) DRB4-dependent TAS3 trans-acting siRNAs control leaf morphology through AGO7. Curr Biol, 16, 927–932.

87. Yifhar, T., Pekker, I., Peled, D., Friedlander, G., Pistunov, A., Sabban, M., Wachsman, G., Alvarez, J.P., Amsellem, Z. and Eshed, Y. (2012) Failure of the tomato trans-acting short interfering RNA program to regulate AUXIN RESPONSE FACTOR3 and ARF4 underlies the wiry leaf syndrome. Plant Cell, 24, 3575–3589.

88. Ding, B., Xia, R., Lin, Q., Gurung, V., Sagawa, J.M., Stanley, L.E., Strobel, M., Diggle, P.K., Meyers, B.C. and Yuan, Y.W. (2020) Developmental Genetics of Corolla Tube Formation: Role of the tasiRNA-ARF Pathway and a Conceptual Model. Plant Cell, 32, 3452–3468.

89. Nagasaki, H., Itoh, J., Hayashi, K., Hibara, K., Satoh-Nagasawa, N., Nosaka, M., Mukouhata, M., Ashikari, M., Kitano, H., Matsuoka, M. et al. (2007) The small interfering RNA production pathway is required for shoot meristem initiation in rice. Proc Natl Acad Sci U S A, 104, 14867–14871.

90. Montgomery, T.A., Howell, M.D., Cuperus, J.T., Li, D., Hansen, J.E., Alexander, A.L., Chapman, E.J., Fahlgren, N., Allen, E. and Carrington, J.C. (2008) Specificity of ARGONAUTE7-miR390 interaction and dual functionality in TAS3 trans-acting siRNA formation. Cell, 133, 128–141.

91. Xia, R., Xu, J. and Meyers, B.C. (2017) The Emergence, Evolution, and Diversification of the miR390-TAS3-ARF Pathway in Land Plants. Plant Cell, 29, 1232–1247.

92. Yu, B., Bi, L., Zhai, J., Agarwal, M., Li, S., Wu, Q., Ding, S.W., Meyers, B.C., Vaucheret, H. and Chen, X. (2010) siRNAs compete with miRNAs for methylation by HEN1 in Arabidopsis. Nucleic Acids Res, 38, 5844–5850.

93. Edgar, R., Domrachev, M. and Lash, A.E. (2002) Gene Expression Omnibus: NCBI gene expression and hybridization array data repository. Nucleic Acids Res, 30, 207–210.

