## Supplementary Figures for "Nuclear and cytoplasmic RNA exosomes and PELOTA1 prevent miRNA-induced secondary siRNA production in Arabidopsis"

#### Figure S1.

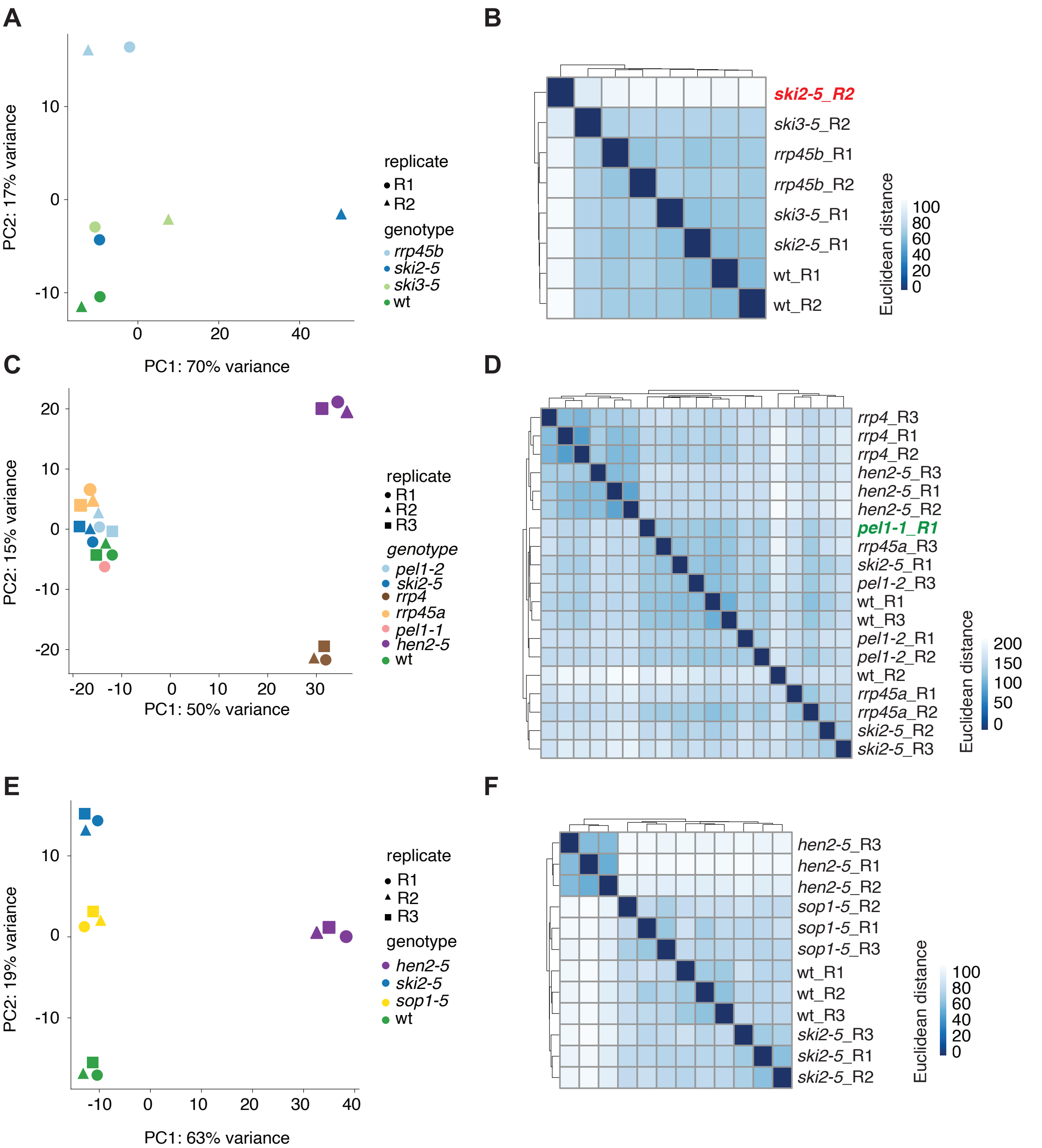


**Supplemental Figure 1. Validation of small RNA sequencing data prior to DESeq analyses.**
**(A)** and **(B)**, small RNA sequencing experiment A, used for analyses reported in Figure 2; **(C)** and **(D)**, small RNA sequencing experiment B, used for analyses reported in Figures 3-7; **(E)** and **(F)**, small RNA sequencing experiment C, used for analyses reported Supplemental Figure 3.
**(A)** Principal component 1 and 2 of WT, *ski2-5, ski3-5* and *rrp45b.* The genotypes are distinguished by color and the two biological replicates are distinguished by shapes. **(B)** A distance matrix of WT, *ski2-5, ski3-5* and *rrp45b*. The second biological replicate of ski2-5 is written in red to indicate that it is an outlier. Based on the principal component analysis and the distance matrix, this sample was excluded from the DESeq analysis.
**(C)** Principal component 1 and 2 of WT, *ski2-5, pel1-1, pel1-2, rrp4, rrp45a* and *hen2-5*. The genotypes are distinguished by color and the biological triplicates in the pool of libraries are distinguished by shapes. **(D)** A distance matrix of WT, *ski2-5, pel1-1, pel1-2, rrp4, rrp45a* and *hen2-5.* The only sample of *pel1-1* is written in green to indicate its inclusion in downstream DESeq analysis as it clusters together with the *pel1-2* samples.
**(E)** Principal component 1 and 2 of WT, *ski2-5, hen2-5* and *sop1-5.* The genotypes are distinguished by color and the biological triplicates in the pool of libraries are distinguished by shapes. **(F)** A distance matrix of WT, *ski2-5, hen2-5* and *sop1-5*.

**Figure S2.**

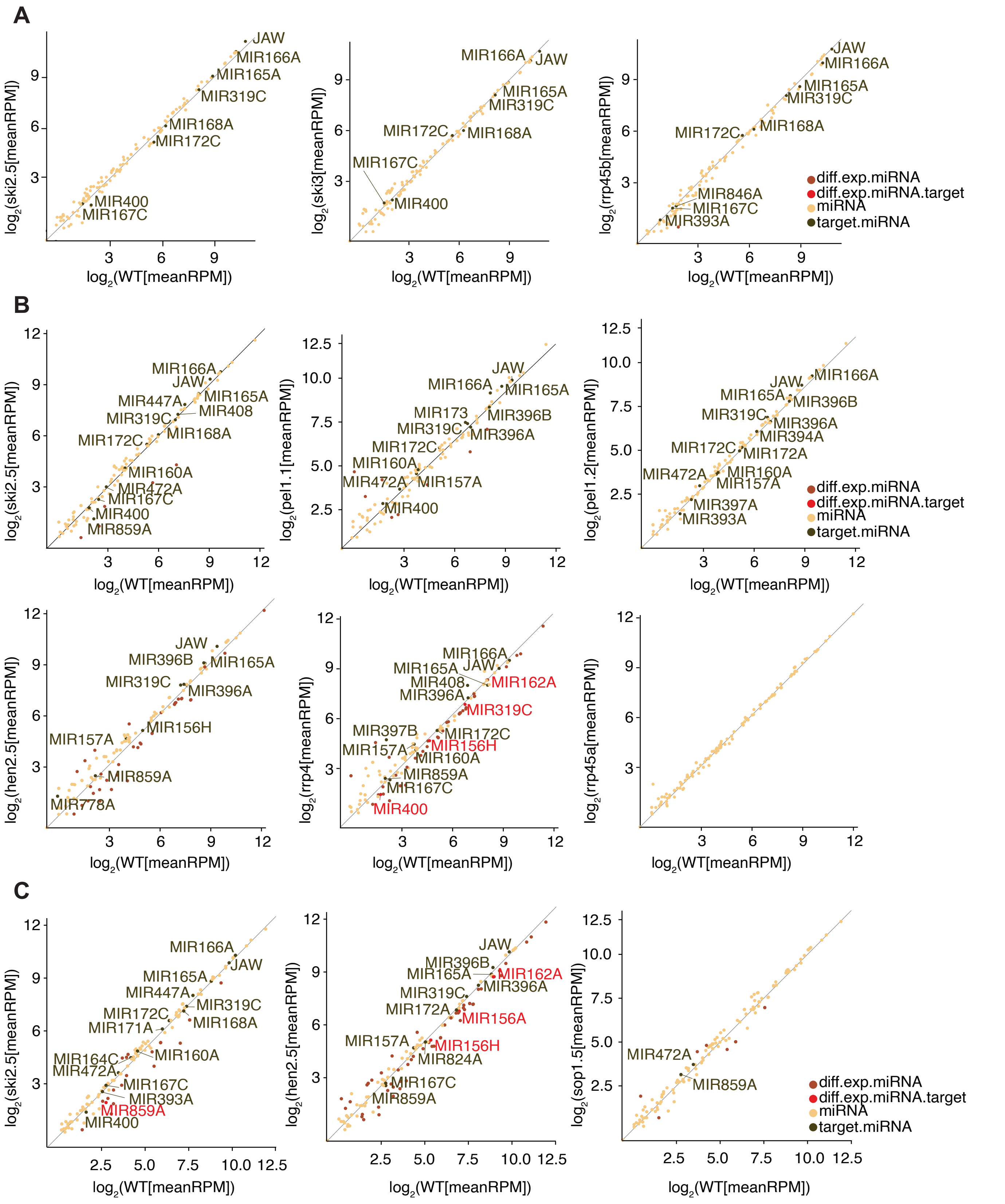


**Supplemental Figure 2. miRNA expression levels in mutants used in this study compared to Col-0 WT.** **(A,B,C)** Scatter plots showing mean RPM of all miRNA genes in mutants compared to Col-0 wild type. Black, miRNA with mRNA targets that give rise to different siRNA levels compared to wild type; yellow, miRNA with mRNA targets with no apparent difference in siRNA levels compared to wild type; brown, differentially expressed miRNA; red, differentially expressed miRNA with mRNA targets that give rise to different siRNA levels compared to wild type. **(A)** Experiment A, miRNAs in *ski2-5, ski3-5* and *rrp45b* compared to Col-0 WT. **(B)** Experiment B, miRNAs in *ski2-5, pel1-1, pel1-2, hen2-5, rrp4* and *rrp45b* compared to Col-0 WT. **(C)** Experiment C, miRNAs in *ski2-5, hen2-5* and *sop1-5* compared to Col-0 WT.

### Figure S3

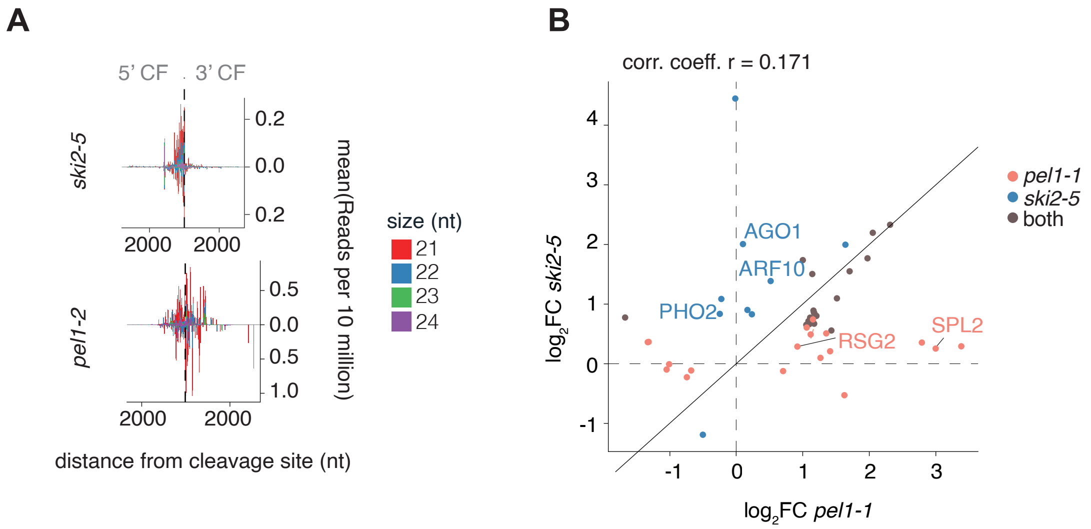


**Supplemental Figure 3. miRNA-triggered siRNA accumulation in *ski2-5* and *pel1* mutants**
**(A)** Metaplot of siRNA read densities (RP10M) along miRNA target transcripts with significantly higher siRNA production in mutants than in wild type. Position 0 is defined by miRNA-guided cleavage sites. **(B)** Scatter plot of log2 of the fold change of read densities (RPM(mutant)/RPM(WT)) of siRNAs (log2 FC) mapped to miRNA targets in *pel1-1* (x-axis) and *ski2-5* (y-axis). Only miRNA targets with significantly higher siRNA read counts in either mutant compared to Col-0 are included. miRNA targets shown in Figure 3B are indicated.

#### Figure S4.

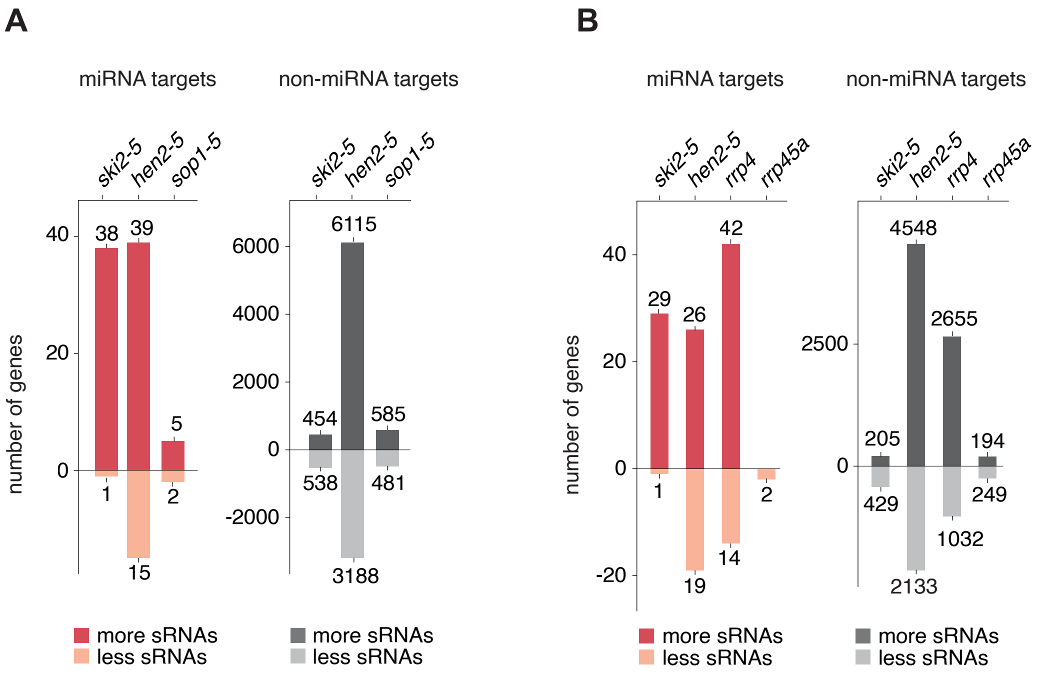


**Supplemental Figure 4. siRNA accumulation in *rrp45a* and *sop1-5* mutants.**
**(A)** Bar plots depicting number of known miRNA targets (red bars) or number of non-miRNA targets (grey bars), which produce either more or less secondary siRNAs in *ski2-5, hen2-5* or *sop1-5* compared to Col-0 WT (Wald test, P < 0.05). The enrichment of miRNA targets in genes producing more siRNAs is highly significant in *ski2-5* (Fisher-test: P <2.2 x 10-16). In contrast, the proportion of miRNA targets found in genes with lower levels of siRNAs in the *ski2-5* compared to WT is not significant (Fisher-test: P = 0.1373). The enrichment of miRNA targets in genes producing more siRNAs is not significant in neither *hen2-5* nor *sop1-5* (Fisher-test for hen2-5: P = 0.2024, Fisher-test for sop1-5: P=0.813). The proportion of miRNA targets found in genes with lower levels of siRNAs in the *hen2-5* and *sop1-5* mutants compared to WT is also not highly significant (Fisher-test for *hen2-5*: P = 0.04279, Fisher-test for *sop1-5*: P= 0,5976). **(B)** Bar plots depicting number of known miRNA targets (red bars) or number of non-miRNA targets (grey bars), which produce either more or less secondary siRNAs in *ski2-5, hen2-5, rrp4-2* or *rrp45a* compared to Col-0 WT (Wald test, P < 0.05). The enrichment of miRNA targets in genes producing more siRNAs is significant in *ski2-5* and *rrp4* (Fisher-test for *ski2-5*: P < 2.2 x 10-16, Fisher-test for *rrp4-2:* P=0.0001719). In contrast, the proportion of miRNA targets found in genes with lower levels of siRNAs in the *ski2-5* and *rrp4* mutants compared to WT is not significant (Fisher-test for *ski2-5*: P = 1.00, Fisher-test for *rrp4:* P= 0.07551). The enrichment of miRNA targets in genes producing more siRNAs is not significant in *hen2-5* and *rrp45a* (Fisher-test for *hen2-5:* P = 0.1223, Fisher-test for *rrp45a*: P=0.4122). This is the same for the proportion of miRNA targets found in genes with lower levels of siRNAs in the *hen2-5* and *rrp45a* mutants compared to WT (Fisher-test for *hen2-5*: P = 0.6169, Fisher-test for *rrp45a*: P= 1).

#### Figure S5.

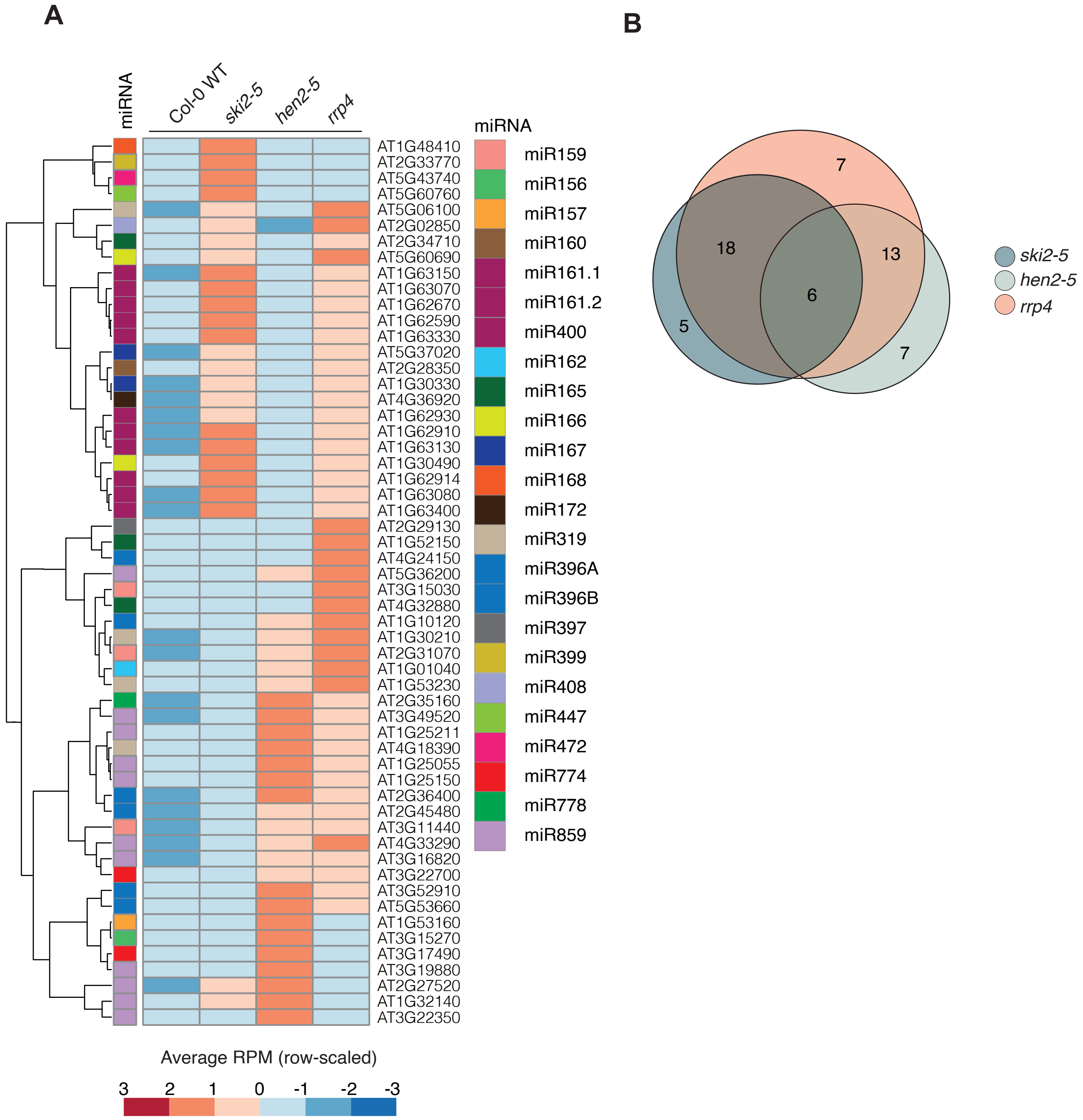


**Supplemental Figure 5. siRNA production from miRNA targets in *ski2-5, hen2-5* and *rrp4-2***
**(A)** Heatmap of known miRNA targets with significantly more siRNAs produced in *ski2-5, hen2-5* and *rrp4-2* compared to WT (padj. < 0.05). In the heatmap, the z-score of the mean RPM of siRNAs mapped to each miRNA target in WT, *ski2-5, hen2-5* and *rrp4-2* are used. The heatmap is clustered by targets. The identity of miRNA targets represented in each is indicated to the right, and miRNA members known to cleave the corresponding targets are indicated with a color to the left of each row. **(B)** Euler diagrams showing overlap in siRNA-producing miRNA targets between *ski2-5, hen2-5* and *rrp4* mutants
