## Supplementary Table S1 for "Nuclear and cytoplasmic RNA exosomes and PELOTA1 prevent miRNA-induced secondary siRNA production in Arabidopsis"

| Oligoname | Sequence (5' to 3') | Purpose |
|---|---|---|
| ski2-5_Salk_LP | GAACTGGTCTTTTTGTCGTGC | Genotyping PCR |
| ski2-5_Salk_RP | TAAATTTGCGGACATTTGAGG | Genotyping PCR |
| ski3-5_GK_LP | AGATGAGGCTTTTGAGAGTT | Genotyping PCR |
| ski3-5_GK_RP | ATTAGCGCATTCCATAACAGATTC | Genotyping PCR |
| ski8-1_Salk_LP | ACAGAGAGACCACGAGAGCAG | Genotyping PCR |
| ski8-1_Salk_RP | GAAGCAAATAAAAACTCCACTGC | Genotyping PCR |
| pel1-1_Sail_LP | GAGAAGCTGTGGAACGAATC | Genotyping PCR |
| pel1-1_Sail_RP | GGCATACCAAGCCCTTAG | Genotyping PCR |
| pel1-2_GK_LP | CGAGTCTGTATGTTCCTTCAC | Genotyping PCR |
| pel1-2_GK_RP | GCATGGAGAACCTCACC | Genotyping PCR |
| rrp45a_GK_LP | CGTGATCATACATCCACCCGAAG | Genotyping PCR |
| rrp45a_GK_RP | ATGCCGCTTCACCTTCTGTG | Genotyping PCR |
| rrp45b(cer7-3)_Sail_LP | CTGGCTGTTCTGGTTGGAGT | Genotyping PCR |
| rrp45b(cer7-3)_Sail_RP | CATTTCCAGAGCCGTTCATT | Genotyping PCR |
| hen2-4_Salk_LP | TATGGTATTCAGCAACCTCCG | Genotyping PCR |
| hen2-4_Salk_RP | GTTCCTCAAATGCTGCTCTTG | Genotyping PCR |
| hen2-5_GK_LP | GACTTGTGAAAGCGCTTTTTG | Genotyping PCR |
| hen2-5_GK_RP | TATGGTATTCAGCAACCTCCG | Genotyping PCR |
| sop1-5_Salk_LP | GGCGAGCAATGAGTTGAATCG | Genotyping PCR |
| sop1-5_Salk_RP | ACTTCGCCAACACCTTATCACC | Genotyping PCR |
| TDNA_GABI08474 | ATAATAACGCTGCGGACATCTACATTTT | Genotyping PCR |
| Salk_TDNA_Lbb1.3 | ATTTTGCCGATTTCGGAAC | Genotyping PCR |
| Sail_TDNA_LB3 | TAGCATCTGAATTTCATAACCAATCTCGATACAC | Genotyping PCR |
| rrp4-2(SOP2)_Eco47I_F | CTATTCCCGTCAACCATGACG | Genotyping PCR |
| rrp4-2(SOP2)_Eco47I_R | CATCGACCTCGGAAGTTCCATGT | Genotyping PCR |
| rdr6-12 BfaI F | TGCAAGAGGAACGTGTGAGGTG | Genotyping PCR |
| rdr6-12 BfaI R | GCTTCAACCTCTTGTACGCATC | Genotyping PCR |
| AGO1_5'_F | AGAGAAGAACGATGCTCCA | probe |
| AGO1_5'_R | TTGTTGCTGTTGTGGTGGTT | probe |
| AGO1_3'_F | GGATTTGCACCATTATGAT | probe |
| AGO1_3'_R | TCAAGAACCTGCAGAGCTT | probe |
| CSD2_5'_F | CCAAACGTCAAACATAGCAGCA | probe |
| CSD2_3'_R | CCGCGGAAACAACTGTCAAC | probe |
| CSD2_5'_F | GGGTGACCTGGGAAACATAA | probe |
| CSD2_3'_R | TCAAGCCAATCACACCACAT | probe |

**SUPPLEMENTARY TABLE S1.** Oligonucleotides used in the study.
